# Mapping the sequence specificity of heterotypic amyloid interactions enables the identification of aggregation modifiers

**DOI:** 10.1101/2021.06.04.447096

**Authors:** Nikolaos Louros, Meine Ramakers, Emiel Michiels, Katerina Konstantoulea, Chiara Morelli, Teresa Garcia, Sam D’Haeyer, Vera Goossens, Dominique Audenaert, Frederic Rousseau, Joost Schymkowitz

## Abstract

Heterotypic amyloid interactions between related protein sequences have been observed in functional and disease amyloids. While sequence homology seems to favour heterotypic amyloid interactions, we have no systematic understanding of the structural rules determining such interactions nor whether they inhibit or facilitate amyloid assembly. Using structure-based thermodynamic calculations and extensive experimental validation, we performed a comprehensive exploration of the defining role of sequence promiscuity in amyloid interactions. Using this knowledge, we demonstrate, using tau as a model system, that predicted cross-interactions driven by sequence homology indeed can modify nucleation, fibril morphology, kinetic assembly and cellular spreading of aggregates. We also find that these heterotypic amyloid interactions can result in the mis-localisation of brain-expressed protein sequences with prevalent activities in neurodegenerative disorders. Our findings suggest a structural mechanism by which the proteomic background can modulate the aggregation propensity of amyloidogenic proteins and discuss how such sequence-specific proteostatic perturbations could contribute to the selective cellular susceptibility of amyloid disease progression.

## Introduction

Neurodegenerative disorders are a diverse group of pathologies that are associated to the gradual deterioration of different brain regions and cause variable clinical phenotypes that range from cognitive impairment to motor deterioration and neuropsychiatric symptoms^1,2^. Despite this complexity, these diseases share fundamental characteristics in regard to their mechanistic underpinnings and clinical manifestation. To begin with, they are characterized by the presence of β-rich amyloid aggregates, the formation of which is initiated by self-propagation of certain proteins and affects particular areas of the brain^3-8^. Another shared clinical feature relates to their specific spatial and temporal progression patterns that can predictably discriminate between distinct disorders by matching symptoms to the functionality of the affected brain regions^9-11^. Efforts to address the basis of cellular and regional vulnerability have focused on the intricate balance between intrinsic neuronal homeostasis to the heterogeneity of amyloid self-assembly and transcellular propagation pathways^9,12^. Genetic variability^13,14^, extrinsic clearance pathways^15^ and molecular expression profiles^16,17^ are important risk factors that enhance cellular susceptibility to toxic amyloid aggregates, with their effects being further exacerbated when coupled to the progressive decline of molecular proteostatic mechanisms that deteriorate with physiological ageing^18^. Although the exact cellular interactions that contribute to the modulation of neuronal susceptibility still remain largely unknown, the prominent role of cellular proteomic heterogeneity in this process is no longer ignored^19^. Specific protein hetero-interactions have been shown to directly influence susceptibility to various amyloid-forming proteins, including among others, Aβ^20-22^, tau^23,24^ and α-synuclein^25-27^ that are involved in Alzheimer’s (AD) and Parkinson’s disease (PD), respectively. In the same line, cell-specific inherent metastability of proteins that supersede their solubility levels has been proposed as a generic mechanism that can promote regional protein co-deposition^28-32^. Cellular predilection to toxic aggregates is also conformation-specific, as recent evidence has shown that different amyloid fibril morphologies derived from the same misfolded protein can characterize other neurodegenerative disorders^33-35^. Regardless of their protein of origin and self-assembly conditions, however, amyloid fibrils share a common structural cross-β architecture^36-39^. Further to this, disease-related amyloid conformers share overlapping thermodynamic distribution profiles, as specific segments that also drive their nucleation predominantly stabilise their amyloid framework^40^. These regions, previously identified as aggregation prone regions (APRs)^41-45^, form thermodynamically stable steric zipper interfaces that staple together amyloid fibril structures. As a result, they are also able to support their own self-assembly^46-50^, as well as to promote heterotypic interactions dominated by sequence similarity^19,51-55^ that have been shown to promote pathology^56-59^ or the formation of biologically functional amyloids^60-66^. Based on the above, here we focused on investigating sequence promiscuity of amyloid core APRs, as a novel structural mechanism that engages in heterotypic amyloid interactions. By performing a comprehensive thermodynamic evaluation of the entire sequence space for APR cores derived from several amyloidogenic proteins, we highlight sequence dependencies that support heterotypic interactions both *in vitro* and within a cellular environment. Together, our results highlight that this novel structural mechanism may be implicated in selective cellular vulnerability by utilising local sequence similarity to promote the entrapment of protein components with important functions, but can also be harvested as the means to improve on high-end therapeutics against major amyloid diseases.

## Results

### Thermodynamic profiling of heterotypic amyloid interactions

Several classes of biomolecules have been found to interact with amyloid fibrils. Glycosaminoglycans, RNA, lipids and rotor dyes, among others, selectively interact with binding pockets or other surface features of amyloid polymorphs^67-70^, while chaperones have been shown to bind to the lateral surface of amyloid fibrils during secondary nucleation or fragmentation^71^. Heterotypic interactions between amyloids and proteins have also been found to modify elongation at the growing tips of amyloids suggesting the existence of cross-seeding in yeast prions^72^, functional amyloids^63,73^ and disease amyloids^48^. While heterotypic amyloid protein interactions have been observed in different model systems, we still have no understanding on the structural rules determining heterotypic amyloid protein interactions beyond the observation that sequence homology favours heterotypic amyloid interactions. Based on our growing insight into amyloid architecture, however, it is becoming evident that amyloid fibril structure is highly ordered and constrained by specific patterns of tightly interlaced side-chains, which are particularly susceptible to minimal variation. For example, even single disease mutations typically induce significant morphological differentiation that often raises barriers of structural incompatibility between strains^74-76^. We now have a heterogeneous and large enough number of amyloid structures to attempt to gain some first insight on the general principles determining hetero-aggregation events. Given the above, we assembled a collection of 83 experimentally determined APR amyloid core structures derived from 18 distinct proteins (**Table 1**), in order to perform a systematic exploration of the potential impact of the incorporation of homologous sequence segments from unrelated proteins into the amyloid core. Using this dataset, we performed detailed thermodynamic profiling of the energies for cross-interaction (i.e. binding of a homologous sequence segment on the growing fibril tip) and elongation (i.e. docking of additional copies of the homologous sequence) against sequence divergence (**Fig. 1a-1c**). We limited our search to single variants of major APRs as: (i) APRs are the kinetic drivers that promote self-assembly of amyloids^41-44^, (ii) individual amyloid polymorphs share energetic profiles in a sense that they depend on APRs as a common framework of high structural stability to counteract longer regions of structural frustration in their core^40^ and (iii) this approach also supports a deeper understanding of potential tendencies, as assignments are performed at a single residue level.

**Figure 1.**
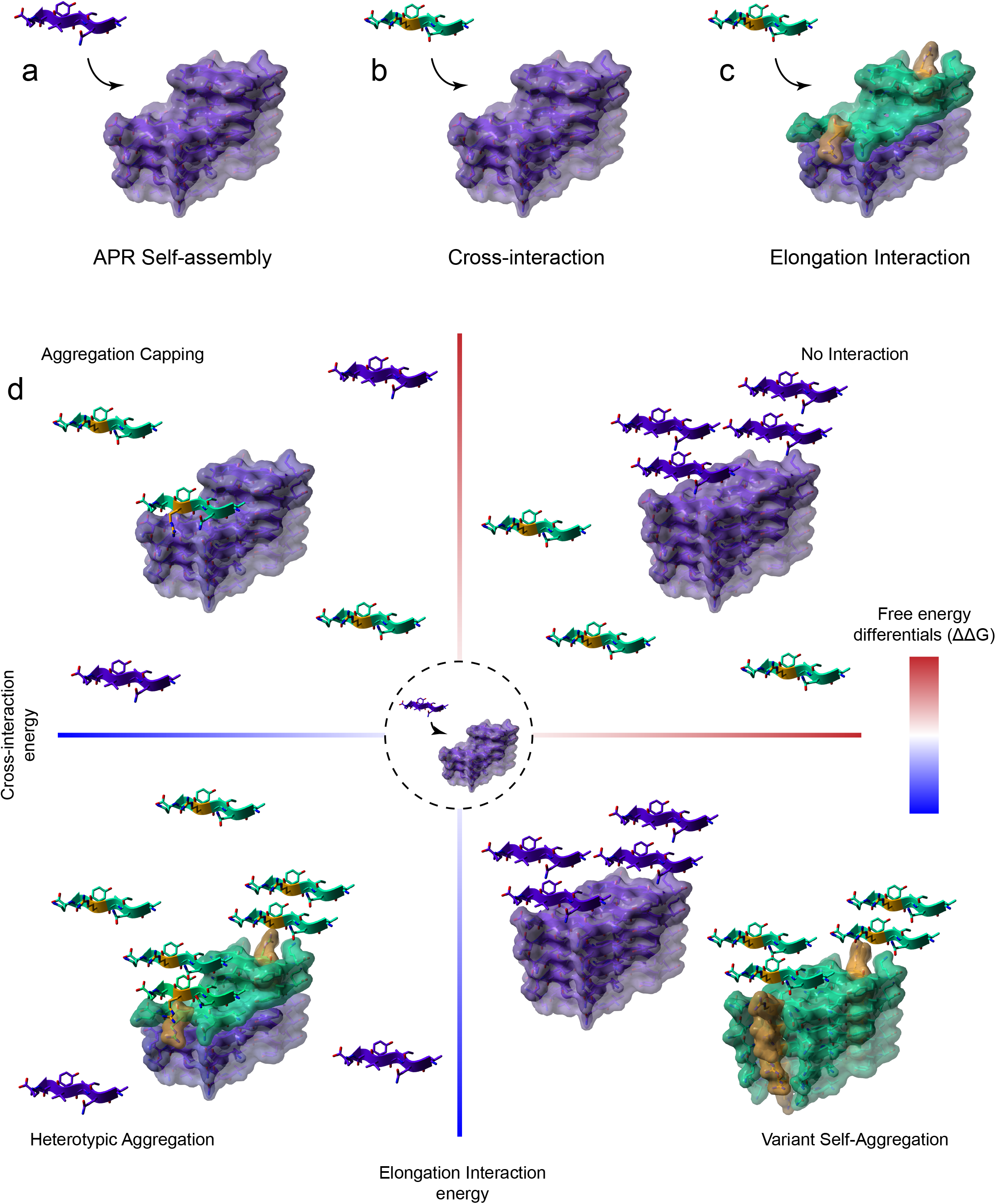
Structural framework to describe cross-interactions of APR aggregation cores. (a-c) Three structural templates were generated for each APR amyloid core structure (shown in purple), corresponding to (a) self-elongation by monomeric APR addition (purple β-strands), (b) primary cross-interaction of a single position (highlighted in yellow) sequence variant (shown as green β-strands) at the amyloid fibril ends and (c) successive elongation by the same variant. (d) Variants that produce favourable differentials compared to monomeric APR elongation are driven towards heterotypic aggregation, compared to disfavourable potentials that limit interactions. Aggregation capping is instigated by sequences that are compatible to cross-interactions with the APR core but block further elongation, while opposite energies are associated to individual self-aggregation, respectively.

We developed a systematic thermodynamic analysis of the impact of side chain mismatches on the cross-interaction and elongation energies in APR cross-beta assemblies, in order to classify homologous sequences into different heterotypic-induced outcomes (**Fig. 1d**). Our approach is based on all-atom structural models of the cross-beta cores formed by the APR regions under study, in which we introduce mismatches and judge their impact on the thermodynamics using the FoldX force field^77^. By comparing the free energy of cross-interaction (**Fig. 1b**) and elongation (**Fig. 1c**) interactions to the free energy potential of the APR self-interaction (**Fig. 1a**), we can define hetero-interaction-compatible variants as sequences that produce thermodynamically favourable cross-interaction free energies at the growing tip of amyloid fibrils. Furthermore, supporting elongation differential energies can distinguish segments that participate in heterotypic assembly (**Fig. 1d**, bottom-left quadrant) from aggregation-blockers (also defined herein as “cappers”) (**Fig. 1d**, top-left quadrant). On the other hand, favourable elongation energies and disfavourable cross-interaction energies suggest a propensity of the variant sequence towards its own intermolecular assembly, thus expected to enhance self-association to cross-aggregation (**Fig. 1d**, bottom-right quadrant). Finally, limited aggregation propensity is expected for stretches that produce unsuited free energy profiles for both modes of interaction (**Fig. 1d**, top-right quadrant).

### Investigating sequence space compatibility of APR cross-interactions

Using our profiling scheme, we investigated structural compatibility for the entire sequence space of single variants of the APR dataset (**Table 1**). Our energetic analysis revealed that less than 1 out of 4 variants engage in both cross-interaction and elongation, i.e. co-aggregation (**Fig. 2a**, bottom-left quadrant), while even fewer sequences (16.9%) were seen to be compatible with suppressing further elongation after cross-reacting with growing fibril ends, i.e. inhibiting aggregation (**Fig. 2a**, top-left quadrant). This apparent incompatibility of APR cores to sequence variation was also supported by the fact that only a limited fraction of sequence variants was predicted to favour self-assembly (6.9%) (**Fig. 2a**, bottom-right quadrant), possibly suggesting that the template backbone arrangements are strongly tailored to their particular sequences. In agreement, increasing the sequence variation to two mismatches further reduced the predicted thermodynamic compatibility, with predictions rendering most homologous stretches containing two mismatches (>75-80%) structurally incompatible for cross-interactions (**Fig. S1**). This is also supported by our previous findings on the cross-reactivity of sequence-targeting engineered anti-viral and anti-bacterial peptide designs^78,79^.

**Figure 2.**
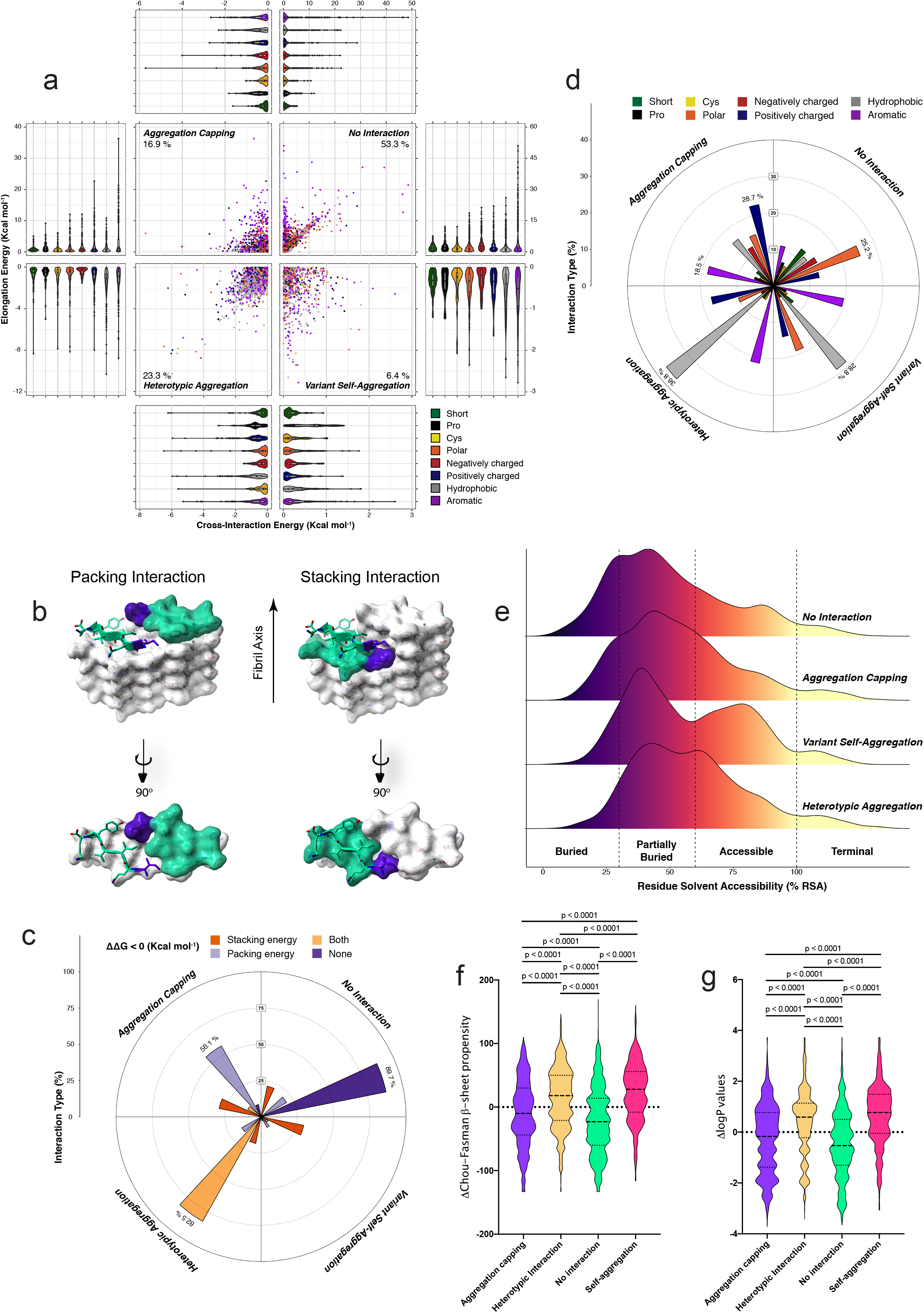
Thermodynamic profiling of APR amyloid core cross-interactions to single variants. (a) Single-mutation potential distributions in the four defined modes of cross-interaction. A quadrant plot is generated by plotting cross-interaction energies on the x-axis and elongation energies on the y-axis, respectively. Energy distributions for each quadrant are shown as violin plots containing box-plots. Residues are categorised as shown in the figure legend (Short=A, G, Pro=P, Cys=C, Polar=N, Q, S, T, Negatively charged=D, E, Positively charged= R, K, H, Hydrophobic=V, I, L, M, Aromatic=F, Y, W). (b-c) Rose plot distribution of the packing and stacking energy contributions along the four modes of interaction. (d) Rose plot distribution of residue type mutations along the four modes of interaction. (e) Residue burial distributions (relative surface area – RSA) for the four modes of interaction. (f) β-propensity and (g) solubility differentials calculated as a difference in value compared to their corresponding cognate APR sequence. Statistics: one-way ANOVA with multiple comparison.

We identified a linear correlation between cross-interaction and elongation free energy potentials for the majority of sequence variants expanding from heterotypic aggregation to non-interactors (**Fig. 2a**). This suggests that selective deviation from this correlation is required in order to develop potent capper designs that efficiently recognise fibril tips and simultaneously disrupt further elongation steps (i.e. strong cross-interaction energies with unfavourable elongation energies). This becomes more evident when comparing cross-sectional packing (residues opposing each other in the same layer) to transversal stacking (residues on top of each in subsequent layers) contributions along the axis of the fibril (**Fig. 2b**). Indeed, almost 90% of unfavourable interactions are described by poor packing and stacking energies, with both interfaces contributing actively for a similar fraction of heterotypic interactors (**Fig. 2c**). Notably, heterotypic capping was found to be facilitated primarily by destabilised packing interfaces during elongation (56.1%), suggesting that although stacking interactions are integral for overall stabilisation, longitudinal packing is more easily destabilised by sequence variation.

Residue distribution analysis pinpointed that hydrophobic side chain substitutions are primarily associated to heterotypic aggregation, however they can also often increase the self-association tendency of variants, leading to independent self-assembly (**Fig. 2a and 2d**). On the other hand, polar side chain substitutions and introduction of so-called gatekeeper residues, such as Pro, Glu and Asp reduce hetero-compatibility, implying that apart of acting as evolutionary suppressors of APRs^80^, these residues may also limit aggregation cross-talk. Besides this, introduction of aromatic and positively charged residues was primarily associated to weak elongation energies. Heterotypic interactions were predominantly associated to partially buried positions, as changes in residues that are tightly packed in the amyloid core were harder to incorporate in co-aggregation compatible variants (**Fig. 2e**). In contrast, high surface exposure reduces specificity and can often increase the self-assembly potential of variants by simultaneously minimising cross-interactions. Finally, β-propensity (**Fig. 2f**) and solubility (**Fig. 2g**) are additional determinants between the four modes of interaction. Mutations that either promoted heterotypic or homotypic assembly were usually associated to increased β-sheet propensity and solubility, compared to their APR counterpart. On the other hand, heterotypic cappers are less soluble and often destabilise β-formation, with the effect being even stronger in the case of non-interacting mutants, respectively.

### Dimensionality reduction reveals the driving forces of heterotypic aggregation

To objectively define the thermodynamic determinants of cross-amyloid interactions, we performed dimensionality reduction and clustering of the individual energy contributions using the Uniform Manifold Approximation and Projection (UMAP) technique (**Fig. 3**). Three primary clusters of potent cappers were identified (**Fig. 3a**, clusters 4 and 6 and to a lesser extent cluster 3) to interact well with fibril tips and significantly disrupt further elongation by mapping cross-interaction and elongation interaction energies (**Fig. 3b**). The first cluster (cluster 4) shows the strongest conversion from stabilising cross-interactions to highly destabilising elongation energies (**Fig. 3b**). Cluster 4 is primarily occupied by aromatics that efficiently cap fibril ends by introducing significant steric clashes during elongation, but not during initial interaction with the fibril tip (**Fig. 3c**). Pure electrostatic repulsion (**Fig. 3d**) can also drive aggregation capping (cluster 3), however is more efficient when coupled with steric hindrance of elongation seen with the longer side chains of the positively charged side chains (cluster 6), but not the negative side chains (cluster 3). Interestingly, globular β-sheet proteins use similar strategies as negative evolutionary invariant designs in their natural folds, in order to prevent uncontrollable edge-to-edge agglomeration^81^, whereas proteins with amyloid-compatible folds, such as β-solenoids, β-rolls and β-ladders are known to be heavily charged, as well as to incorporate polyprolines or aromatic bulges as edge-capping mechanisms to prevent aggregation events at the tip of their folds^82-84^. Another mode of capping refers to disruption of the hydrogen bond network that staples β-strands together in growing amyloids (**Fig. 3e**). This cluster (cluster 1), is in principle mostly composed of proline variants that act as β-breakers. However, this capping mode is less efficient due to the fact that the levels of disruption are thermodynamically low and similar between cross-interaction and elongation (**Fig. 3b**). Finally, short side chains can also mildly cap fibril ends (cluster 2) by gradually weakening the free energy gaining from dispersive interactions between the solute and solvent (**Fig. 3f**), whereas polar and hydrophobic residues are poor cappers that are not particularly driven by specific interactions (cluster 5).

**Figure 3.**
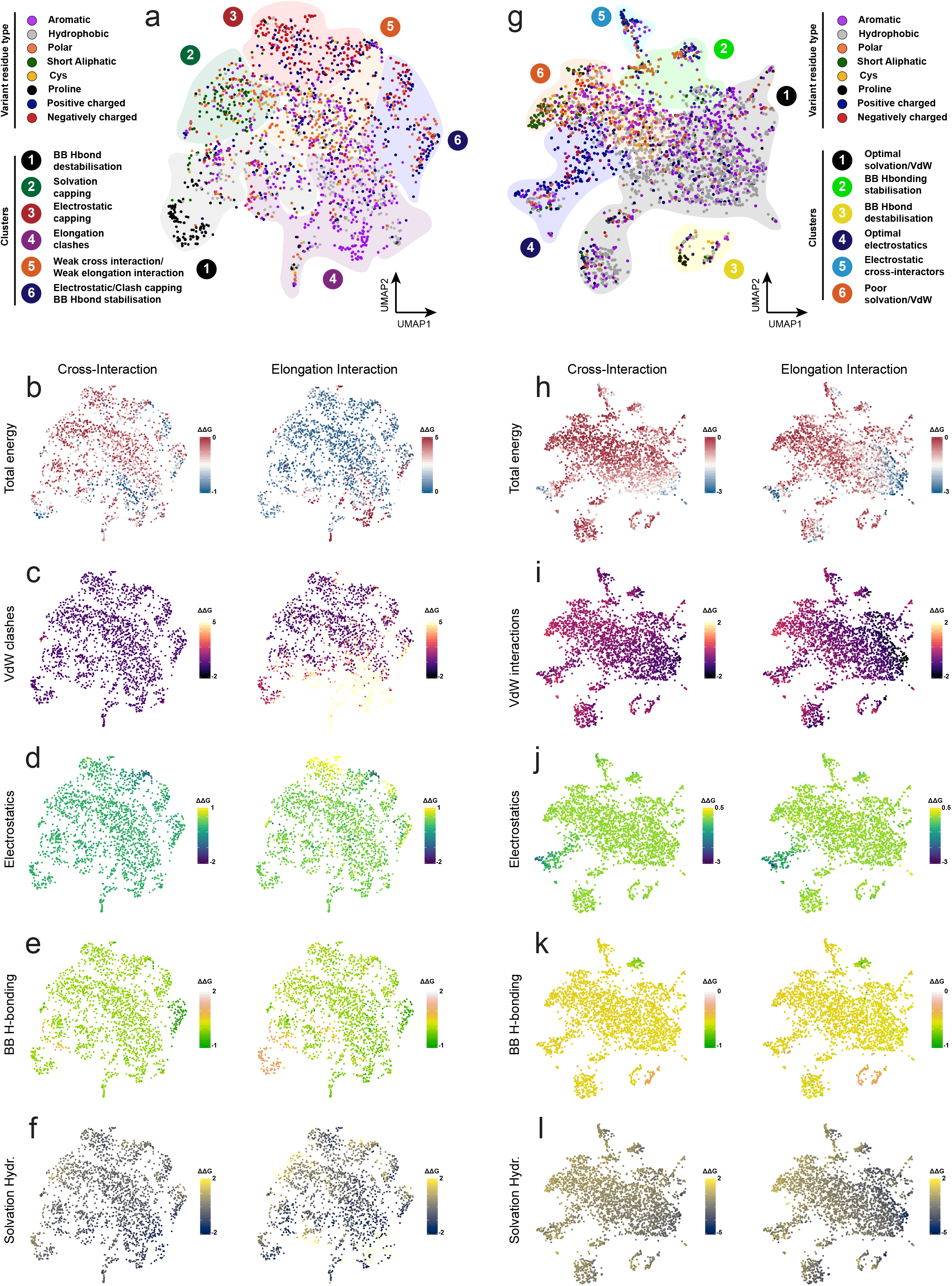
Dimensionality-reduction mapping of the heterotypic sequence space. (a) Clustered modes of interaction that dominate capping variants were identified by analysing (b) total energy and individual energy components, including (c) steric clashes, (d) electrostatics, (e) backbone hydrogen bonding and (f) solvation energy of hydrophobics. (g) Clustered modes of interaction that dominate variants supporting heterotypic aggregation. Independent clusters were identified by analysing (h) total energy and individual energy components, including (i) van der Waals interactions, (j) electrostatics, (k) backbone hydrogen bonding and (l) solvation energy of hydrophobics.

Dimensionality reduction charted a different energy landscape for co-aggregating sequences (**Fig. 3g-3l**). In this analysis, cluster 1 contained the strongest cross-interacting variants (**Fig. 3g**). Composed principally by hydrophobic (and to a lesser extent, aromatic) side chains, this cluster is defined by tightly packed hydrophobic cores that maximise Van der Waals (**Fig. 3i**) and solvation (**Fig. 3l**) contributions and is located at opposite ends of the cross-aggregating sequence space, compared to short and polar side chains (cluster 6). Other cases indicated that electrostatic interactions (**Fig. 3j**) can also stabilise cross-aggregation (cluster 5), however are more potent when further stabilised by optimal side chain stacking during elongation (cluster 4). Finally, backbone hydrogen bonding is a much more limited factor in co-aggregation (cluster 2) and can be rather destabilising when introduced by strong β-breakers, such as Pro residues (cluster 3) (**Fig. 3k**).

### Self-assembly of aggregation prone regions is modified by sequence stretches sharing high sequence similarity

Next, we sought to experimentally investigate these different modes of fibril-tip interactions. For this, we selected a well-known and thoroughly described APR from tau as a case study^85^. The VQIVYK (PHF6) stretch, located in the C-terminal repeat domain of tau, has been demonstrated to be crucial for tau aggregation^86,87^ and is a dominant stabiliser of all tau amyloid polymorphs^40^ (**Fig. 4a**). Thermodynamic profiling of single variants indicated that cross-interacting variants of the VQIVYK sequence occur primarily at partially buried positions, in contrast to substitutions of the fully buried Ile residue that introduce significant steric clashes during cross-interactions, as well as the Lys side chain that has minimal selectivity due to its high solvent exposure (**Fig. 4b**). In line with our UMAP clustering analysis, strong capping variants relied on elongation clashes introduced by aromatic packing (**Fig. 4c**) or on charge repulsion introduced by successive stacking of charged residues during elongation (**Fig. 4d**). On the other hand, strong co-aggregating variants enabled a tighter packing of the hydrophobic core (**Fig. 4e**), utilised electrostatic interactions that promote cross-interactions without causing significant disruptions during elongation (**Fig. 4f**) or better-defined stacking interactions along the surface of the growing fibril core (**Fig. 4g**).

**Figure 4.**
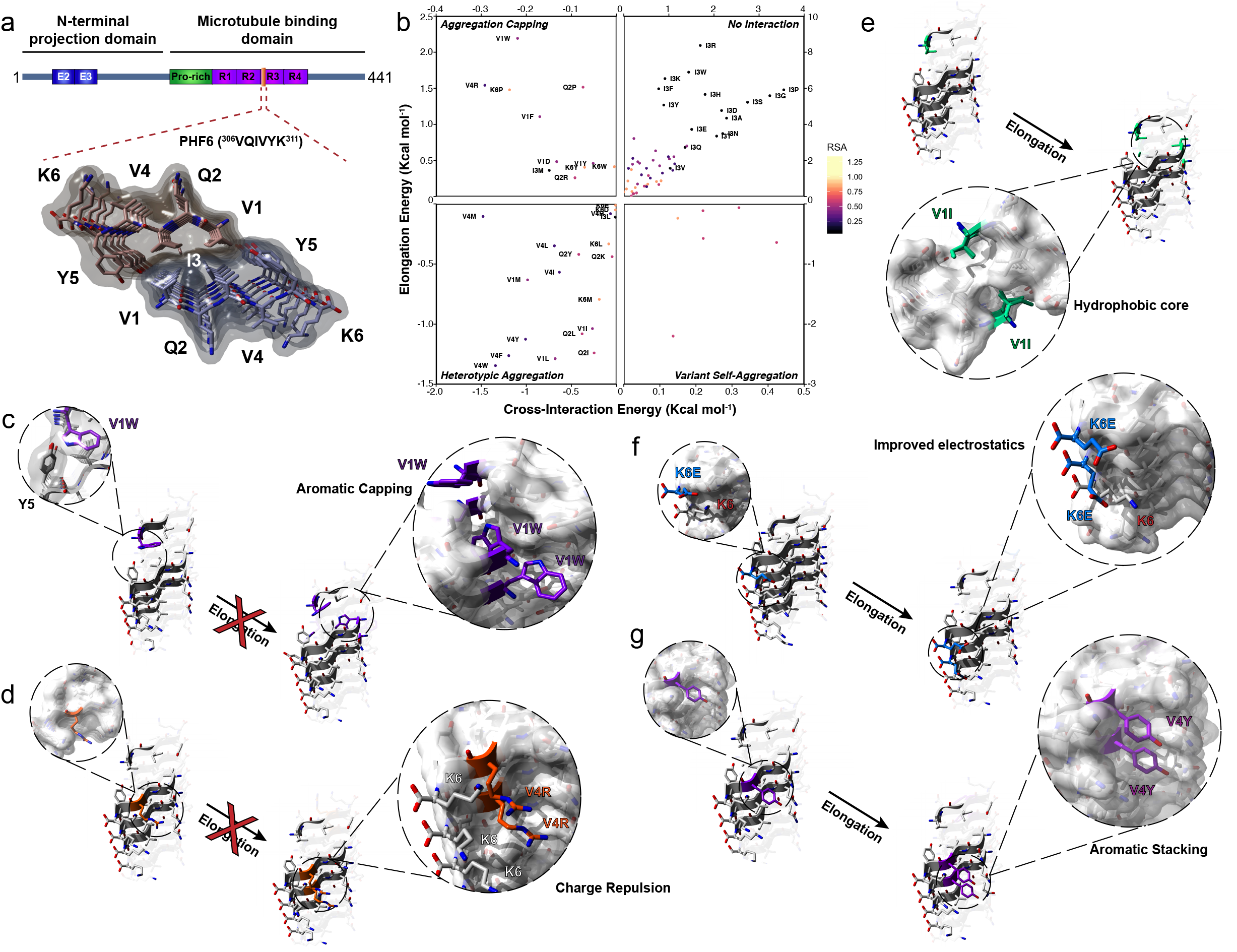
Thermodynamic profiling of cross-interactions for the VQIVYK amyloid core. (a) Schematic representation and positioning of the VQIVYK APR in full length tau. (b) Quadrant plot analysis of the four modes of interactions for single variants of the VQIVYK APR. (c-d) Capping interactions are facilitated by the incorporation of (c) bulky aromatic residues that blocked further elongation at the fibril tips through steric clashes or (d) charged side chains that blocked elongation through electrostatic repulsion of stacked charges. (e-g) Heterotypic aggregation is promoted by (e) hydrophobic mutations that stabilise the aggregation core, (f) electrostatic interactions that improve surface solubility or (g) improved stacking interactions at the exposed fibril surface.

To experimentally investigate these calculations, we synthesized a library of 90 peptides corresponding to 78% of all single amino acid substitutions of the VQIVYK APR. We excluded introduction of cysteines to avoid further complexity introduced by the formation of intermolecular disulphides and also avoided substitutions of the Tyr residue at position five, since it enables fast and accurate readout of peptide concentration. First, peptide:APR mixtures were monitored using Th-T aggregation kinetic assays. For this screen, we used sub-stoichiometric mixtures of the variant peptides against PHF6 (1:5 analogy). The reasoning behind this was that it enabled tracing of subtle differentiation in aggregation kinetics, while at the same time reduced the propensity of most variants to participate in self-assembly. In total, 7 variants, all corresponding to the exposed Lys position were found to still self-aggregate at 25μM and as a result were excluded from further analysis (**Fig. S2**). For the rest, following curve fitting of the monitored Th-T curves (**Fig. 5a, 5b and Fig. S2**), we calculated and summarised fold changes of aggregation half-times (t^1/2^) in a volcano plot, with the negative logarithm of the p-values represented on the vertical axis (**Fig. 5c**). Remarkably, we observed a significant overlap to their thermodynamic profiling, as calculated capping (**Fig. 5c**, green points) and inducing modifier sequences (**Fig. 5c**, purple points) overlapped to peptides that had a negative or positive impact on the experimentally determined aggregation kinetics. Additionally, most variants of the buried central Ile position did not engage in cross-interplay, as seen by the minimal changes in aggregation kinetics (**Fig. 5c**, yellow points). End-state fluorescence analysis validated that most co-aggregating variants increased aggregation (V1I, Q2I, K6E and K6D have reduced Th-T levels, but significantly reduce the kinetic lag-phase) (**Fig. 5d**), whereas diminished Th-T levels supported the inhibitory effect of the capping sequences (**Fig. 5e**). Equilibrium thermodynamics analysis using critical concentration determination showed that co-assembly variants, such as V1I, V4I, K6E, K6D and K6L significantly reduced the concentration of the wild type APR that remains in solution when equilibrium is reached, a clear sign that the free energy of aggregation was impacted (**Fig. 5f**). Conversely, aromatic, charged and proline substitutions effectively capped and reduced aggregation of VQIVYK, as up to a five-fold increase of the wild type APR was identified in the soluble fraction of those mixtures, even after a week of incubation (**Fig. 5g**). The latter was also confirmed using electron microscopy, since almost no amyloid formation was observed for V1W, K6P, V1Y and V1F mixtures at the same timeframe (**Fig. 5h**).

**Figure 5.**
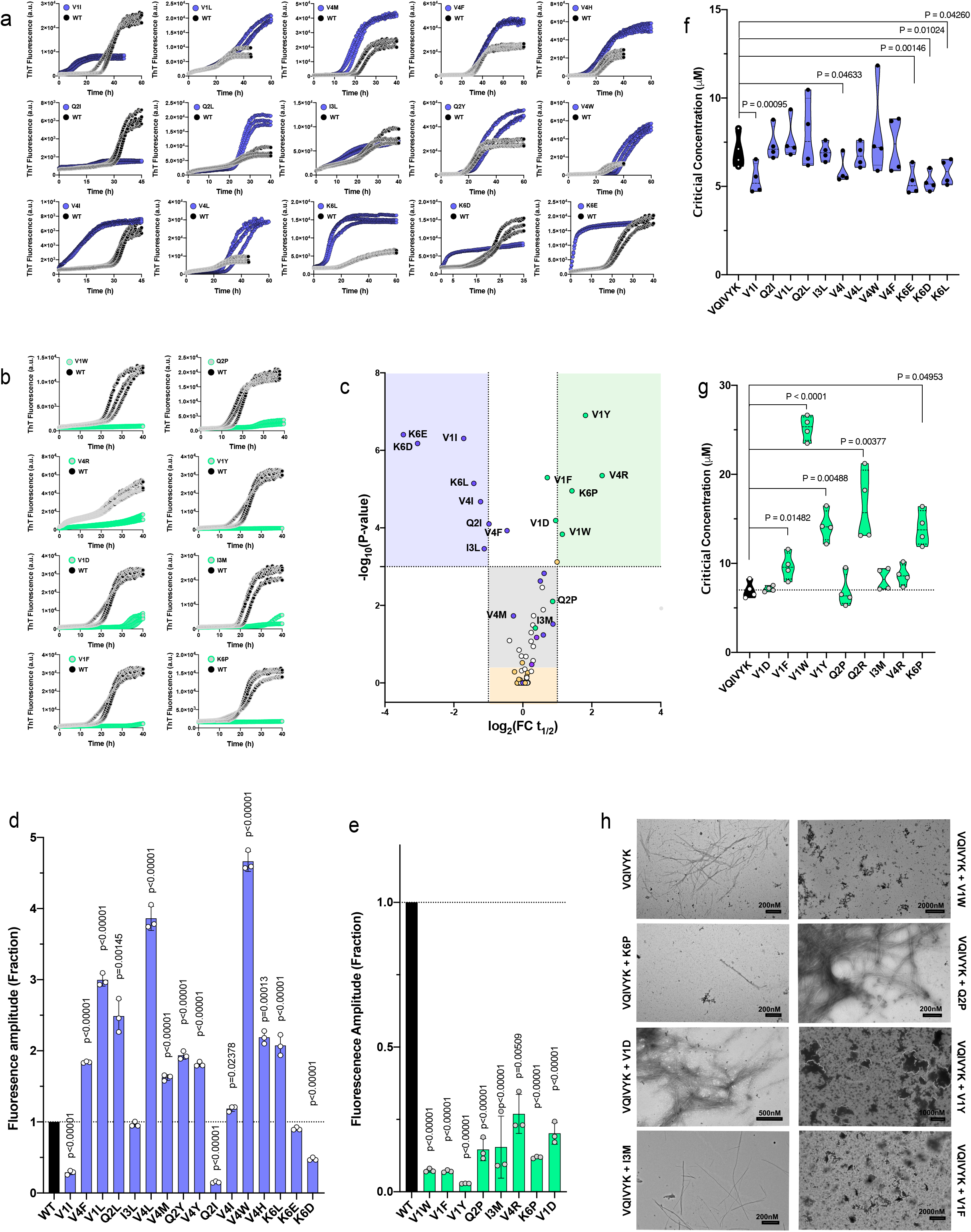
Peptide screening of variant cross-interactions with the VQIVYK aggregation prone region from tau. Th-T kinetic assays (n=3) of VQIVYK-alone (125μM) or in the sub-stoichiometric presence of the strongest (a) co-aggregating or (b) inhibiting single position variants (25μM). (c) Volcano plot analysis of the kinetic halftimes for the entire peptide screen (Fig. S3). Green-and blue-shaded backgrounds indicate capping and co-aggregating sequences of high significance. Sequences with a strong thermodynamic profile for capping or heterotypic interaction are shown in green or purple points, respectively, whereas mutants of the Ile residue are shown in yellow. Statistical significance was determined using Tukey’s test for multiple comparisons (compared to VQIVYK-alone halftime). (d-e) End-state fluorescence and (f-g) critical concentration modifications induced by the strongest (d-f) heterotypic aggregating and (e-g) capping sequences. Statistical significance was determine using Tukey’s test for multiple comparisons (compared to VQIVYK-alone). (h) Electron micrographs of capping mix samples after 7 days of incubation. Minimal to no fibril formation was observed for the strongest cappers (V1W, K6P, V1Y, V1F).

To further ensure the findings of the UMAP clustering, we also analysed the co-aggregation kinetics of a smaller subset of single variants derived from another experimentally defined APR segment from human Apolipoprotein A-I (ApoA-I)^88,89^. Using the same thermodynamic profiling against a model of the ApoA-I APR, we randomly selected 5 of the strongest hetero-aggregating and capping variant sequences (**Fig. S4a-b**). Following peptide synthesis, our experimental observations once more supported the heterotypic profiling, since Th-T screening followed by kinetic and end-state analysis (**Fig. S3c-e**) indicated that all 10 variants had an expected modulatory effect on the aggregation kinetics of the WT sequence.

### Sequence-dependent modifiers alter the morphology of APR amyloid fibrils

Previous studies have indicated that even single mutations can have notable effects on the morphology of amyloid fibrils^74-76,90-96^. Therefore, we employed transmission electron microscopy to investigate if the substoichiometric presence of heterologous APRs, such as described in the previous paragraph, could also alter the morphology of fibrils formed by the VQIVYK APR. Mixtures of conserved variants, such as V1I and V4I, produced longer and thicker fibril networks compared to cognate APR self-assembly. On the other hand, mixtures containing co-interactors incorporating more radical mutations that contain charge inversions, such as K6E or K6D, caused significant morphological differentiation, by forming super-twisted helical fibrils with very tight pitch distances, while the K6L variant produced fibril fragments of shorter lengths (**Fig. 6a**). To gain further insight on this conformational heterogeneity, we used fluorescence probe binding. Due to its excellent sensitivity, this approach has been used in past studies to determine structural heterogeneity of fibril populations derived even from the same protein constituent^35,97^. Fluorescence spectral acquisitions were obtained side-by-side by adding pFTAA (**Fig. 6b**) or curcumin (**Fig. 6c**) to fibrils obtained from peptide mixtures. In each case, the derived spectra were cross-compared to those produced by peptide-only solutions, as well as against solutions containing APR-only amyloid fibrils. Spectral analysis of the pFTAA and curcumin aggregation reporters indicated the presence of different amyloid conformers represented by spectral shifts of band maxima, as well as from inter-band ratio variations. To increase the discriminative sensitivity of the reporters, we coupled this approach to principal component analysis (PCA). Towards this, we normalised the derived spectra after background subtraction and fed the resulting points to PCA. We found that this way structural conformers were actively separated, as the primary principal components (PCs) accounted for more than 90% of the variability in both dye spectra (**Fig. 6d and 6e**). The eigen space defined by pFTAA spectral analysis resulted in almost complete separation between different conformers (**Fig. 6d**). More specifically, the charge switch of the exposed Lys in the case of mixtures containing K6D and K6E resulted in the formation of equally distant (from PHF6 fibrils, cluster 1), yet closely-related diversified amyloid polymorphs. This is evident by the fact that both the peptide-alone (clusters 9, 11) and co-assembly samples (clusters 8,10) gathered close together in the eigen space defined both by pFTAA and curcumin spectra (**Fig. 6e**), as well as from their observed shared super-twisting morphology. Similarly, the K6L mixture polymorphs (cluster 12) were equally distant to WT amyloid fibrils in terms of their pFTAA binding capacity. Hydrophobic variants also formed distinct conformers that were clearly defined with pFTAA, and to a less extent with curcumin (clusters 2, 4, 6), however their reduced eigen distances to the PHF6 cluster (cluster 1) suggest that these differentiated fibril polymorphs are more similar to the APR fibrils, at least in terms of their dye-binding surface properties.

**Figure 6.**
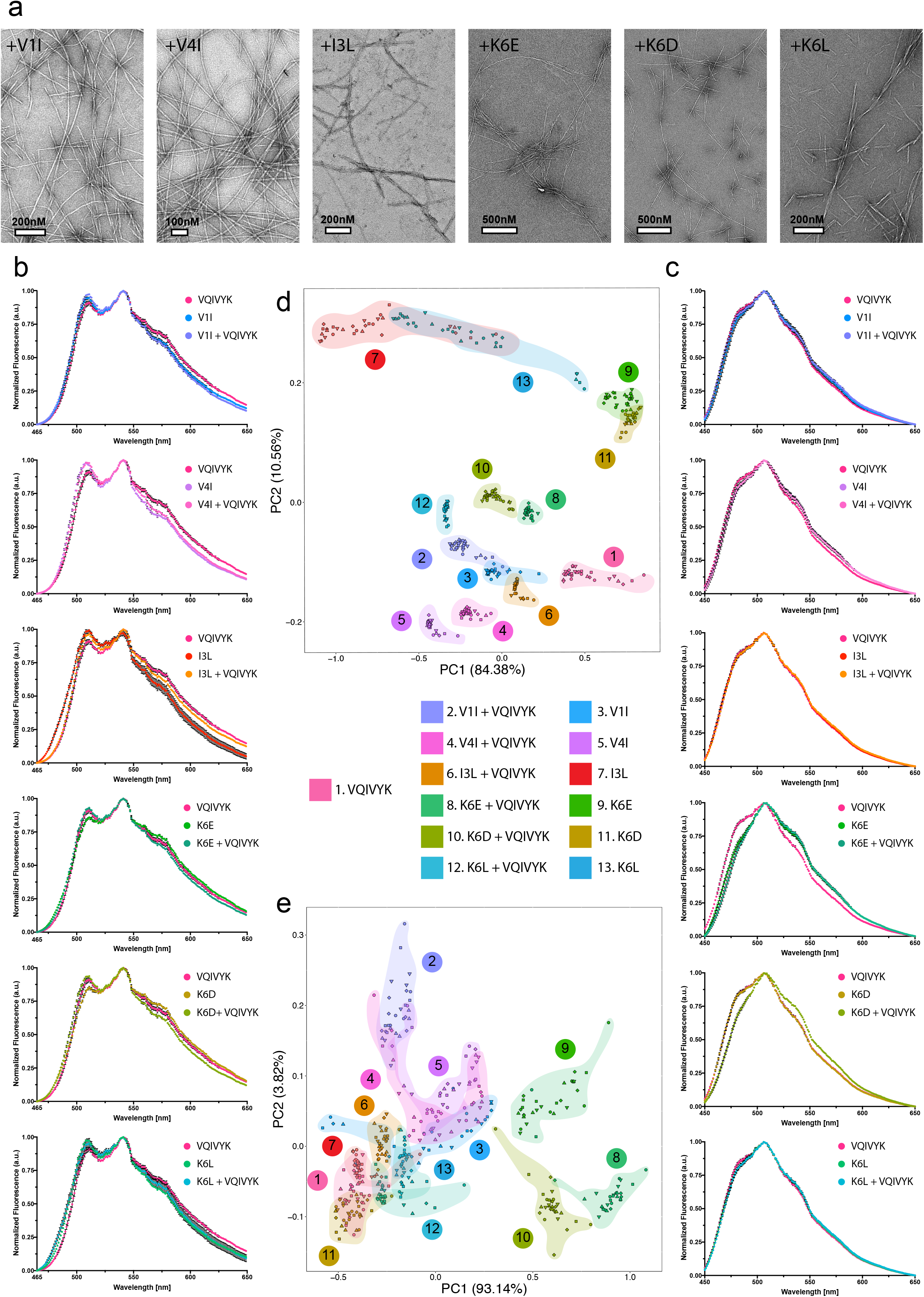
Morphological differentiation induced by co-aggregating sequence-dependent variants of the VQIVYK aggregation prone peptide. (a) Electron micrographs indicate that co-aggregating sequences modify the morphology of amyloid fibrils formed compared to the VQIVYK-alone fibrils. (b-c) Normalised binding spectra of (b) pFTAA and (c) curcumin amplified from fibrils derived from mix, peptide modifier-alone or VQIVYK-alone samples. (d-e) PCA of the derived (d) pFTAA and (e) curcumin spectra highlighted the distribution of heterogenic conformers that cluster in the defined eigen space. For each sample, six individual preparations were split in five independent aliquots and combined in thirty data points (n=30) per sample in order to represent the intrinsic variability in the fluorescence measurements.

As an additional manner to independently validate the morphological differentiation of the fibrils formed in the mixtures, we employed Fourier-Transform infrared spectroscopy (FTIR) coupled to PCA analysis. This approach supports the identification of structural divergence in amyloid polymorphs by applying vibrational spectroscopy directly on amyloid fibrils and as such, is independent of the physicochemical properties of external probes. Amyloid fibril samples produce strong peaks in the amide I and amide II regions (wavenumber region 1500-1700 cm^-1^), mainly arising from the stretching and bending vibrations of carbonyl-and NH groups, respectively, that hold together the β-backbones that constitute their axis. As a result, we isolated this spectral region from each sample (**Fig. 7a**), normalised and fed the resulting points to PCA. Once again, the derived eigen space distributions indicated that mixtures containing variants with modified charge content form strains that cluster in close proximity (clusters 8, 10 and 9, 11) (**Fig. 7b**). Importantly, however, FTIR spectral analysis highlighted that mixtures containing the more conserved hydrophobic variants also produce fibrils (clusters 2, 4, 6) that are structurally different from WT amyloid fibrils (cluster 1), despite the fact that they closely resemble the dye binding properties of the latter, suggesting that they may share similar exposed fibril surfaces that facilitate dye binding but ultimately form structurally distinct aggregation cores.

**Figure 7.**
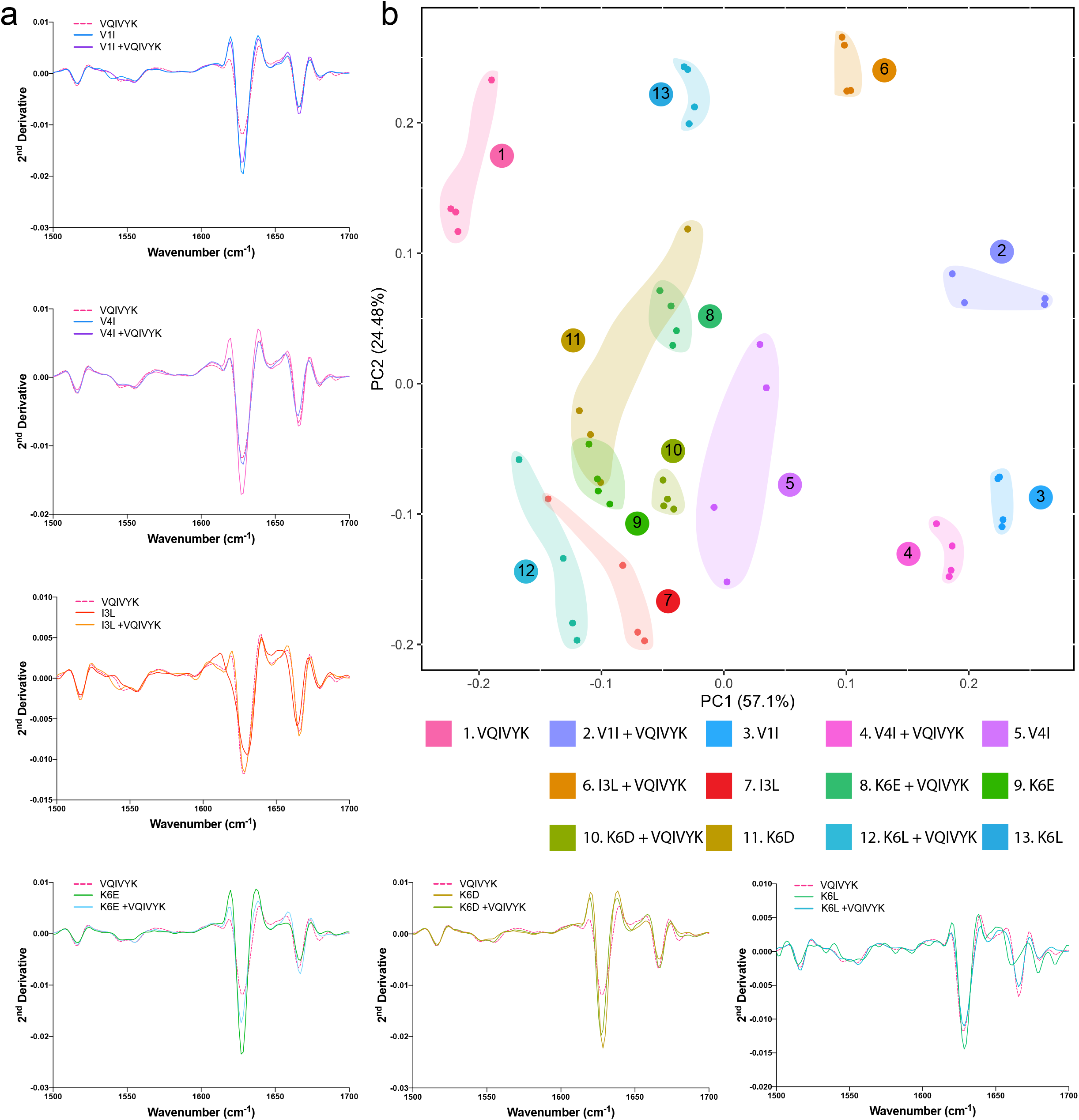
FTIR spectroscopy coupled to PCA revealed the formation of diversified conformers in the presence of VQIVYK co-aggregating modifiers. (a) Second derivatives of the FTIR spectra generated from mixed, peptide modifier-alone or VQIVYK-alone samples, focused around the amide I and amide II region (1700cm^-1^-1500cm^-1^). (b) PCA analysis of the derived spectra indicates the presence of different clustering locations in the eigen space representing the formation of differentiated amyloid fibril conformers in the different samples. For each sample, two individual preparations were split in two independent aliquots and combined in four data points (n=4) per sample in order to represent the intrinsic variability in the spectral measurements.

### Over-expression of full-length proteins harbouring APR variants modulates aggregation in cells

Our *in vitro* screening showed that short sequence stretches with homology to APRs are potential aggregation modifiers, supporting accumulating data on sequence-driven amyloid cross-interactions^19,51-54^. To extend this notion further, we sought to assess whether full-length proteins harbouring such homologous hotspots are vulnerable to cross-aggregation. Based on a proteome-wide search for PHF6 sequence homologs and subsequent manual curation, we selected and tested a subset of 11 full length and/or domain regions of proteins containing such co-aggregation hotspots, in addition to full-length (tau^2N4R^) and repeat-domain tau (Tau^RD^) which were included as controls (**Table 2**). In order to experimentally investigate if these protein regions can indeed participate in cross-talk and modulate tau aggregation, we designed constructs for transient expression (**Fig. 8a**). To distinguish expressing from non-expressing cells, each gene construct included a fluorescent reporter (mKO2), separated by an internal ribosome entry site (IRES). The constructs were transfected into HEK293 Tau^RD^-P301S-CFP/YFP expressing biosensor cells that are highly sensitive reporters of tau-specific seeding-competent aggregates^98^. Recombinant full-length tau aggregation was monitored *in vitro* (**Fig. S5a**) and used to produce uniform tau seeds by sonicating end-state fibrils (**Fig. S5b**) that were then concentration-dependently transfected into biosensor cells (**Fig. 8a**). This yields a concentration-related gradient induction of aggregation of the cellular tau reporter construct that can be quantified through image analysis by counting the formation of FRET-positive puncta. Using this experimental setup, we compared the seeding capacity of exogenously added tau aggregates in cells expressing our constructs compared to controls, and we verified the aggregated nature of the resulting cellular inclusions, using fluorescence recovery after photobleaching (**Fig. 8b and Fig. S6**). Moreover, construct colocalization with the tau inclusions was traced using immunofluorescence (HA-staining) (**Fig. 8c**). High-content screening revealed that six of the selected constructs colocalize with tau FRET-positive inclusions (**Fig. 8c**, merged channel). Cells that strongly expressed these constructs were significantly more susceptible to seeding of tau aggregation, as induced tau aggregation raised by at least 20% in transfection-positive cells, when compared to both non-expressing, as well as to non-transfected cells (vehicle control) and even increased to 30% for specific constructs at high seeding concentrations (IDE, TRA2B and DOCK3) (**Fig. 8d**). Impressively, concentration-dependent quantification analysis revealed that this effect remained even when treating with lower concentrations of seeds, with certain proteins (**Fig. 8c**, IDE and TRA2B) rendering cells vulnerable to tau seeding even at picomolar concentrations, whereas no visible aggregation was observed at similar conditions in the corresponding controls. On the other hand, over-expression of tau^RD^ and tau^2N4R^, using the same construct design, did not have any effect on the efficiency of tau aggregation and spreading in the cells, which was also recapitulated for the rest of the constructs included in the selected subset (**Fig. S7a**).

**Figure 8.**
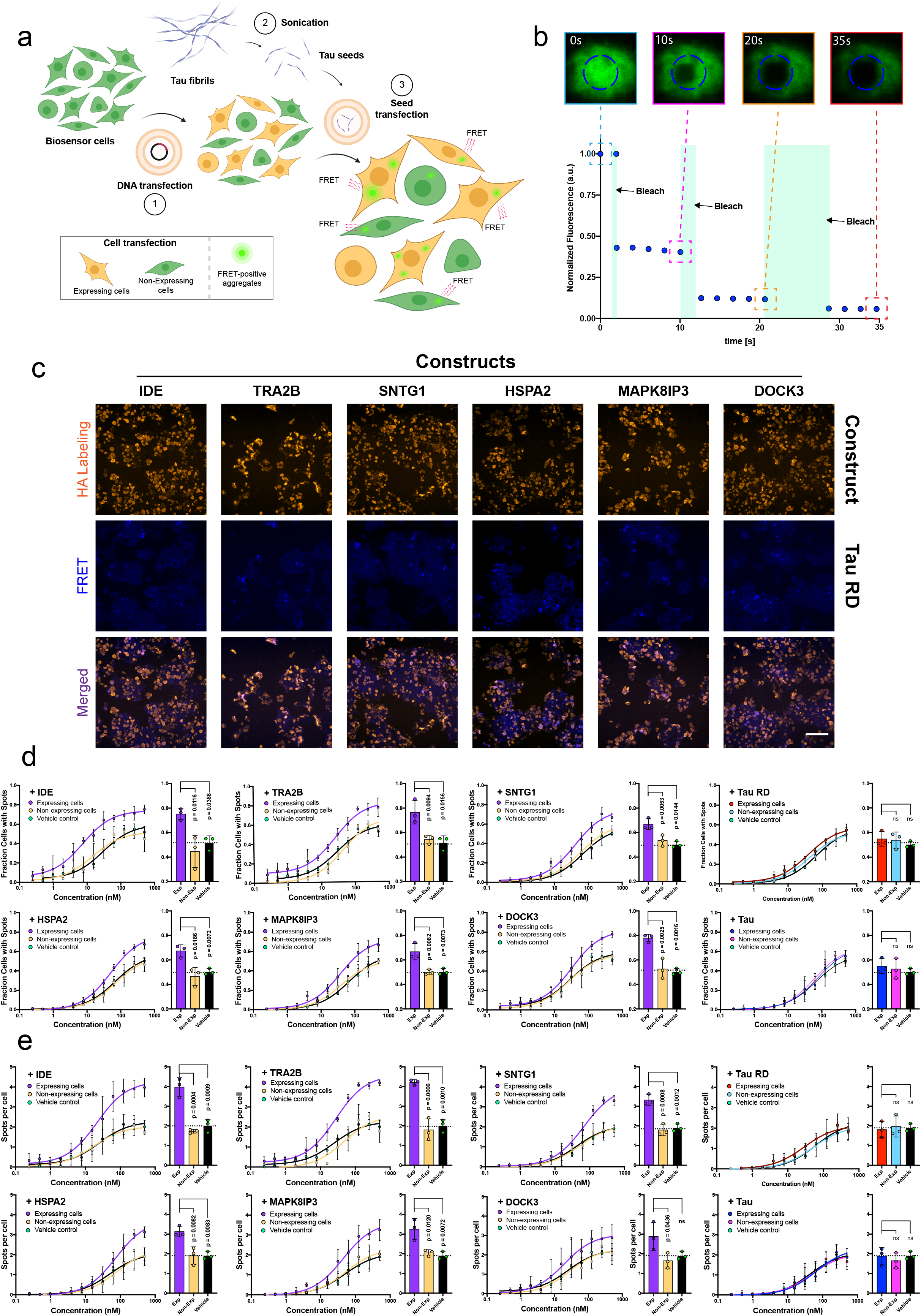
Proteins harbouring localised sequence promiscuity to the VQIVYK aggregation prone peptide modify susceptibility to tau spreading in FRET biosensor cells. (a) Graphical depiction of the experimental setup in the tau biosensor cells. Transient expression of protein constructs, followed-up by secondary tau seeds transfection is measured by quantifying the formation of individual FRET-intensive puncta in construct-expressing (traced with HA-staining) and non-expressing cells. (b) FRAP measurements of FRET-intensive puncta in the biosensor cells. Complete absence of fluorescence recovery was observed after every successive bleaching step of fluorescence puncta. (c) Representative images of cells expressing individual constructs (HA staining channel) containing tau inclusions shown as fluorescent puncta (FRET channel). Merging of the two channels indicates significant colocalization (purple regions) between HA-intense and FRET-intense regions in expressing cells (Bar = 100μm). (d-e) Absolute quantification of (d) the number of construct-expressing and non-expressing cells containing tau aggregates (n= 3 independent experiments) or (e) the number of spots per cell after dose-dependent treatment with tau seeds, compared side-by-side to the vehicle control (no construct transfection), as well as to tau^RD^ and 2N4R transfected cells, respectively. Bar plots highlight differentials observed in cells when treated with the highest concentration of tau seeds (500nM). Statistical significance was calculated using one-way ANOVA with multiple comparisons.

Another indication that the transient over-expression of these constructs increases cellular susceptibility to tau aggregation was highlighted by the morphological analysis of cellular inclusions (**Fig. 8e**). Results showed that with the exception of DOCK3, there was a concentration-dependent increase in the number of inclusions formed per cell, with certain constructs having a doubling or higher effect (IDE, TRA2B and SNTG1) when exposed at higher concentrations of tau seeds, compared to the controls. Similarly, no significant morphological differentiation was observed when transfecting with either tau^2N4R^, tau^RD^ or any of negative constructs (**Fig. S7b**), respectively, thus further supporting the notion that apart of simply participating in co-aggregation with cellular tau inclusions, these proteins may also actively enhance cellular susceptibility to tau aggregation.

### Optimising the design of structure-based amyloid inhibitors

Recent developments have pointed out that sequence-driven structured-based inhibition of amyloids can be an effective approach to counter amyloid formation^55,87,99-107^. In agreement, our thermodynamic profiling and umap reduction analysis also revealed that certain modes of interaction are more successful in capping the ends of growing aggregates, highlighting that aromatic variants have the strongest potential by introducing steric hindrance during elongation. We validated this notion *in vitro* by showing that strong cappers of the VQIVYK APR also often incorporated aromatic residues, while the V1W capper specifically also reduced amyloid formation and critical concentration after several days of co-incubation. Coupling this approach to the recent burst of cryo-EM structures of different tau strains^34^, the V1W capper is also expected to be primarily efficient against recombinant tau amyloid fibrils (as well as to strains related to CBD or Pick’s disease), due to the central position of the PHF6 segment in their amyloid core (**Fig. 9a**). Previous studies have also proposed that modified scaffolds designed to maximise interaction (e.g. tandem or microcyclic designs) and impose structural constraints can enhance the activity of structure-based inhibitors^106,108^, a notion that was also validated during our previous work on antiviral, antibacterial and cancer-cell targeting aggregation-prone peptide designs^78,79,109^. Following this premise, we designed a tandem peptide (named CAP1) incorporating the V1W capping sequence and experimentally tested its capacity to inhibit tau aggregate formation. *In vitro* Th-T kinetics validated the potency of the CAP1 capping activity, as it successfully inhibited the self-assembly of both the PHF6 hexapeptide (**Fig. 9b**) and recombinant full-length tau (tau^2N4R^) (**Fig. 9c**). To investigate the targeting specificity of CAP1 towards aggregate species of tau, we utilized microscale thermophoresis (MST). Towards this end, we generated fluorescently labelled tau seeds by sonicating end-state amyloid fibrils formed after co-incubation of ATTO_633_-labelled and unlabelled tau (1:9 analogy). Dose-response affinity analysis disclosed that CAP1 specifically binds to tau seed aggregates with high affinity (EC_50_ = 145 ± 49 nM), whereas no significant binding was observed against monomeric tau, respectively (**Fig. 9d**). Similar to this, seeding inhibition was also calculated in the biosensor cell line by counting the formation of FRET-positive spots as a function of CAP1 concentration. The derived dose-response curve revealed a high inhibitory effect for CAP1 with an impressive IC_50_ of about 200 nM (**Fig. 9e and 9f**), that is very similar to the determined binding affinity of the peptide and corresponds to a five-fold or higher improvement in efficacy compared to optimal tau inhibitors from previous studies^87^. Overall, our results highlight that due to the current surge in amyloid template structures^39^, our growing structural knowledge of amyloids constitutes thermodynamic profiling, coupled to optimized scaffold design, a competent strategy to design novel aggregation suppressors of high-specificity.

**Figure 9.**
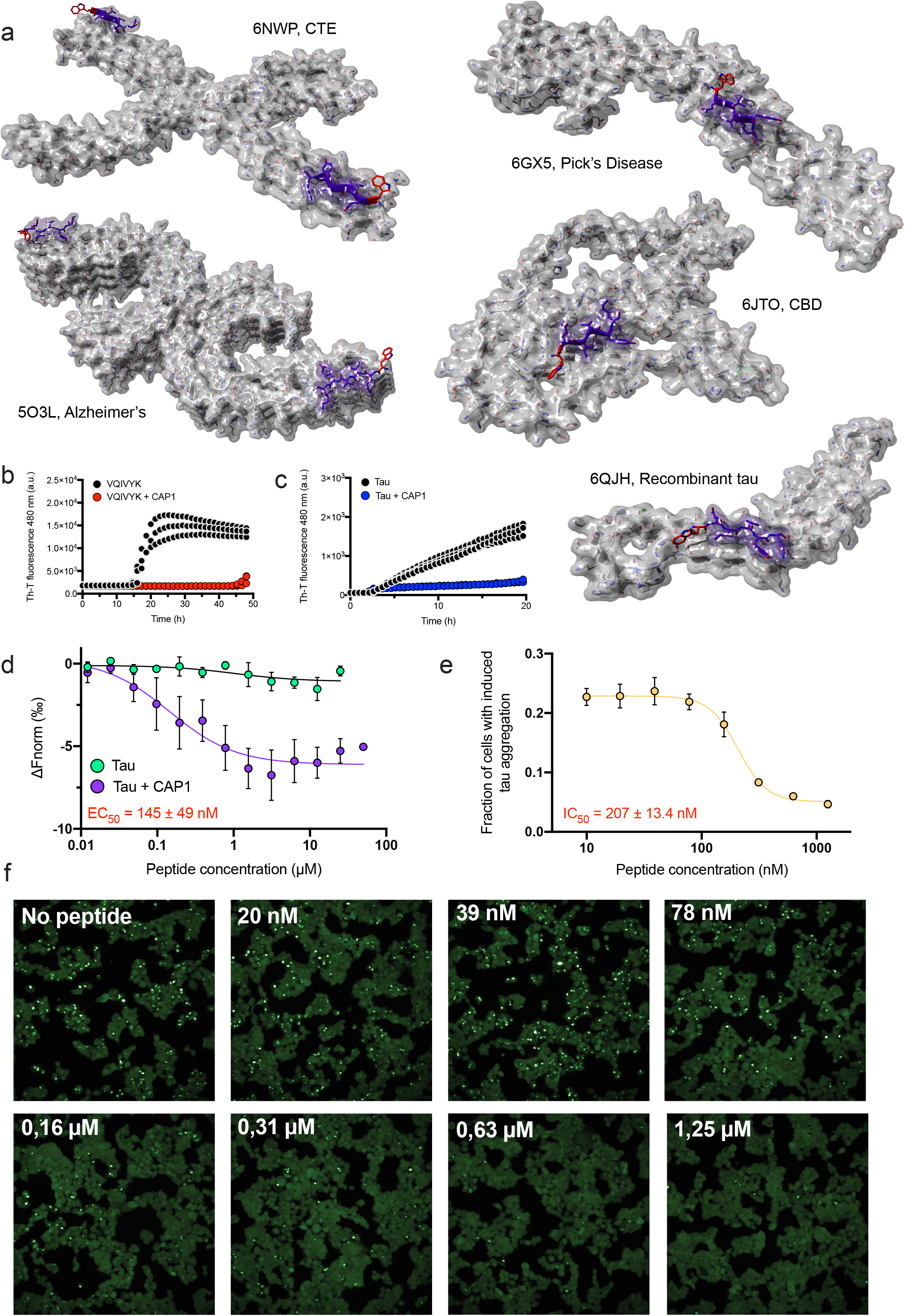
Structure-based inhibition of tau aggregation. (a) Structure-based design of a thermodynamic strong VQIVYK-targeting capper variant sequence (V1W), using full length tau fibril polymorph cryo-EM structures. (b-c) The CAP1 peptide inhibits both (b) VQIVYK and (c) full-length tau aggregation, as shown by Th-T aggregation kinetics. (d) High affinity binding of CAP1 to tau aggregation seeds (purple curve, 145 ± 49nM). No binding was determined for CAP1 to soluble monomeric tau (green curve), suggesting a high specificity of aggregated species. (e-f) Dose-dependent inhibition of tau seeding in the FRET biosensor cell line after pre-treatment of tau seeds (125nM) with incremental concentrations of the CAP1 peptide. (e) An inhibitory concentration value (IC_50_) of 207nM was determined using curve fitting analysis. (f) Representative images of biosensor cells treated with tau seeds pre-incubated with incremental dosage of the CAP1 peptide. Higher CAP1 concentrations significantly reduce the formation of tau inclusions shown as FRET-intensive puncta.

## Discussion

Recent developments in structure determination methodologies, such as cryo-EM, microcrystal electron diffraction and solid-state NMR have provided significant advancement in the field of amyloids. Our structural insight on different architectures of amyloid polymorphs, APR aggregation cores and even oligomeric species is now reaching levels that support a broader understanding of the key structural features that mediate major amyloid-related properties, such as their self-assembly mechanisms, kinetics and overall structural stability^39-50^. On the other hand, our knowledge on amyloid cross-talk with other protein components still remains limited. Despite this, more and more evidence is coming to light indicating that cross-aggregation could be on the basis of defining the apparent selective vulnerability of specific cell types to aggregates or complex spatiotemporal spreading patterns of amyloid deposition, while may also explain observed overlaps between distinct pathologies or why certain amyloid conformers are associated to them, respectively^19^. Building on the above, we provided here, for the first time, a deeper understanding of the structural determinants that define sequence dependency of amyloid cross-aggregation interactions. To achieve this, we performed a systematic thermodynamic evaluation, coupled with multidimensionality analysis, to identify the dominant forces that mediate cross-talk with experimentally determined APR amyloid cores. Our results indicated that even for highly conserved sequences, such as single position variants, a thermodynamically favourable fit within the defined aggregation core is rather hardly accommodated. This notion comes to add to our recent thermodynamic profiling of the fibril cores of full-length amyloid fibrils, which highlighted that these APR segments provide an extremely conserved framework that commonly stabilises different polymorphs. Furthermore, the same analysis revealed that although additional segments of the polypeptide chain participate in hetero-packing when incorporated in the fibril core, these segments are described by energetically degenerative tertiary packing^40^, thus supporting our findings on the limitations of cross-aggregation interactions within amyloid cores.

Owing to the above, we next tested whether proteins containing homologous sequence stretches as potential co-aggregation hotspots could be particularly susceptible to the aggregation propensity of amyloidogenic proteins. Our cellular screening assay, using tau as a case study, validated this premise, yet importantly also indicated that these proteins can further influence the seeding efficiency, morphology and spreading of tau aggregates in the cells. These results suggest that sequence-specific modulatory effects can work in parallel to other mechanisms, as for instance supersaturated sub-proteomes^28-32^ or heterotypic-induced biomolecular condensation^110-113^, to influence amyloid interplay with the background proteomic content of various cell types, thus promoting selective cellular vulnerability. This becomes more evident when considering the role of the six proteins that were here found to significantly modify tau spreading in cells, as they have major impact in progression of AD and various other neurodegenerative disorders. In more detail, the insulin degrading enzyme (IDE) is imperative during clearance of Aβ peptide fragments and has been recently designated as a prime target for new therapeutic treatments against both AD and T2D, respectively^114-116^. Our results showed that IDE colocalises in tau inclusions and promotes spreading, processes that precede Aβ accumulation and plaque formation, suggesting that the latter can be amplified by its early entrapment and gradual loss of function in tau aggregates. Similarly, the dedicator of cytokinesis, DOCK3 (also known as modifier of cell adhesion -MOCA and presenilin-binding protein -PBP), is another important protein involved AD progression and several other neurological deficiencies, including tauopathies and CJD. This enzyme is a known interactor of presenilin, a genetic marker involved in AD, and has also been shown to redistribute and accumulate in neurofibrillary tangles extracted from AD brain samples^117^, indicating that it also colocalises in vivo with tau aggregates. Transformer-2 homolog beta (TRA2B) is a splicing factor that controls alternative splicing of the MAPT gene encoding expression of tau. The reportedly altered expression and activity of TRA2B has been directly implicated to major neurological disorders, such as AD and PD, as well as to promoting tau hyperphosphorylation^118-120^. Importantly, this comes to add to recent evidence indicating that several nuclear speckle components, such as TRA2B, mislocalize to cytosolic tau aggregates in cells, mouse brains, and brains of individuals with AD, frontotemporal dementia (FTD), and corticobasal degeneration (CBD)^121^. Synaptotagmin-1 (SNTG1) is essential for proper synaptic transmission and cognitive function. Recent mass spectrometry assays on cerebrospinal fluid extracted form AD patients, highlighted its use as a novel biomarker in dementia^122^. Furthermore, SNTG1 has a compensatory protective function by gradually increasing its binding to presenilin in the aging brain, an association that has been shown to deter in sporadic AD brains^123,124^. The above suggest that gradual depletion of SNTG1 due to co-aggregation with tau can have detrimental cascading effects during AD progression. The MAPK8IP3 gene encodes for JIP3, a neuronally enriched critical regulator of axonal lysosome abundance^125^. Loss of JIP3 functionality in pluripotent stem cells (iPSCs) results in the aberrant accumulation of Aβ42^126^, suggesting that its inactivation by co-aggregation with tau at the early stages of AD brains can be an important initiator for Aβ proliferation. Finally, despite the known activity of Hsp70 in preventing or inhibiting tau aggregation, our assay revealed that a fragment containing both the nucleotide and substrate binding domain of the molecular chaperone is also vulnerable to tau co-assembly, suggesting that as the proteostatic control mechanisms of cells erode over ageing, protective components such as chaperons may worsen the load produced by amyloids. At this point, our work here presents evidence on a novel generic structural mechanism that cultivates sequence-driven interactions of amyloids to various cellular protein components. Future work is required in order to contextualise this structural mechanism to other generic modes that promote heterotypic aggregation or to understand how and if sequence-specific heterotypic knock-down of certain proteins is amenable to spatiotemporal spreading patterns and selective cellular toxicity of neurodegenerative diseases.

Structured-based designs have been used for years as a strategy for the development of new molecular inhibitors in conformational diseases^55,87,99-107^. Following this logic, we also showed here that the accumulating numbers of amyloid structures, combined to detailed thermodynamic profiling of sequence-specific heterotypic interactions can be used to optimise the design of aggregation cappers. By applying this approach on cryo-EM structures of tau polymorphs, we tested the efficacy of a tandem peptide design, CAP1, in blocking tau aggregation *in vitro* and in cells. Our results indicated that CAP1 selectively binds with high affinity to tau aggregates and blocks its cellular spreading with a five-fold improved efficacy compared to previous designs, suggesting that this approach can indeed be a promising methodology for the development of novel therapeutics in amyloidosis diseases.

## Material and Methods

### Thermodynamic profiling using the FoldX energy force field

We collected a complete set of recently published APR amyloid core structures from the PDB^127^ (**Table 1**). First, we utilised the FoldX energy force field^77^ to generate cross-interaction and elongation instances of cross-assembly for every template by mutating single residues of chains located at its fibril ends (**Fig. 1a-1c**). Second, we used FoldX to perform a thermodynamic breakdown of the energy potentials for both modes of interaction. FoldX as a method has been described in length previously^77^, but briefly here, during free energy calculations, the force field first calculates the free energy contribution of each atom in protein interfaces based on its own position relative to neighbours in the complex. Following this, FoldX subsequently sums individual contributions together, first at the residue level, to calculate segment interaction potentials. This allows to accurately chart the free energy contribution (ΔG) of each residue participating in intermolecular interfaces but also reports on individual thermodynamic components (e.g. Van der Waals, electrostatics, H-bonding or electrostatics, entropy) contributing to overall structural stability. Based on this premise, interaction energies per variant were represented as differentials cross-compared to the free energy potential of the wild-type interaction:

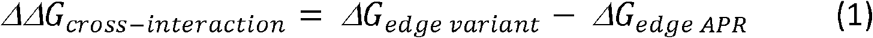

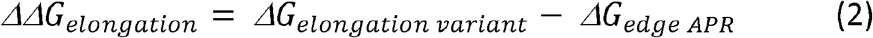

where Δ*G*_*edge variant*_ is the free energy of the cross-interaction of a variant chain at the APR fibril end (**Fig. 1b**), Δ *G*_*elongation variant*_ is the free energy of interaction between a single variant chain docked against an APR axial end occupied by variant chains (**Fig. 1c**) and Δ *G*_*edge APR*_ corresponds to the interaction energy between the cognate APR chain against its own amyloid core (**Fig. 1a**). The reasoning behind using differential ΔΔG values is two-fold: (i) the calculated differentials are comparisons to thermodynamically stable interacting chains derived from experimentally determined APR crystal structures, (ii) while as differentials, they enable global analysis since they only report on the effects in free energy imposed by single mutations and are indifferent to the relevant starting stability of the template structure.

### Uniform manifold approximation and projection (UMAP) dimensionality reduction

A defined sequence space was constructed by merging the identified single-variant capping and cross-aggregating sequences for the complete set of 83 experimentally APR amyloid core structures from 18 proteins (**Table 1**). A 30-dimensional vector, composed by a wide list of individual energy components, including H-bonding, electrostatics, entropy, solvation and Van der Waals interactions between both backbone and side chain atoms, among more, was extracted using the FoldX force field. First, this multidimensional vector was analysed using principal component analysis and the derived principal components were subsequently fed into a umap matrix. Finally, each data point, representing a single-position variant, was reduced and embedded in 2D-space using the R umap package, with the minimum distance to the nearest neighbour set to 0.3 and the number of neighbours to 15, in order to avoid extreme local clustering complexity.

### Peptide library synthesis

Peptides were synthesised using an Intavis Multipep RSi solid phase peptide synthesis robot. Peptide purity (>90%) was evaluated using RP-HPLC purification protocols and peptides were stored as ether precipitates (−20⍰°C). Peptide samples were initially pre-treated with 1,1,1,3,3,3-hexafluoro-isopropanol (HFIP) (Merck), then dissolved in traces of dimethyl sulfoxide (DMSO) (Merck) (<5 %) and filtered through 0.2μm filters before dissolving in the final buffer.

### Aggregation assays

For Th-T kinetics, each peptide variant was pre-treated to form films. The cognate APR peptide was then dissolved and filtered in DMSO, then split into equal aliquots that were used to dissolve the variant films. The resulting mixtures were subsequently dissolved in PBS. Final concentration of the WT APR was set to 125μM and 25μM for the variants (1:5 analogy). Thioflavin-T (Sigma) was added in half-area black 96-well microplates (Corning, USA) at a final concentration of 25μM. Fluorescence was measured in replicates (n⍰=⍰3) using a PolarStar Optima and a FluoStar Omega plate reader (BMG Labtech, Germany) at 30°C, equipped with an excitation filter at 440nm and emission filter at 490nm. To determine kinetic rates, derived spectra were normalised and fitted following:

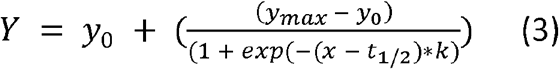

where fluorescence intensity (Y) is represented as a function of time (x). y_max_ and y_0_ indicate maximum and starting fluorescence values, respectively, whereas t_1/2_ and k are the kinetic half times and elongation rates of the fitted curves. t_1/2_ were determined separately for each individual replicate per sample. For endpoint solubility analysis, following incubation for 7 days, peptide mixture preparations were subjected to ultracentrifugation at 76.000 g for 1h at 4oC. The isolated supernatant was mixed with 6M Guanidine-HCl and 0.2% acetic acid and injected into an analytical HPLC. Peptide concentration was then calculating by integrating the AUC values of the peak corresponding to the WT APR peptide.

### Transmission electron microscopy

Peptide mixtures were incubated for 7 days at room temperature in order to form mature amyloid-like fibrils. Suspensions (5μL) of each peptide solution were added on 400-mesh carbon-coated copper grids (Agar Scientific Ltd., England), following a glow-discharging step of 30s to improve sample adsorption. Grids were washed with milli-Q water and negatively stained using uranyl acetate (2% w/v in milli-Q water). Grids were examined with a JEM-1400 120⍰kV transmission electron microscope (JEOL, Japan), operated at 80⍰keV.

### Fluorescence dye binding

For fluorescence dye binding, we prepared equimolar mixtures, variant-only and PHF6 APR-only preparations at a concentration of 500μM in milli-Q water. For statistical analysis, six individual preparations were each split into five aliquots, resulting in thirty in total replicates per sample (n=30) that were left at ambient conditions for seven days to form amyloid fibrils. Suspensions (20μL) of peptide solutions were then mixed with pFTAA and curcumin at 0.5μM and 5μM final concentration, respectively. Fluorescence emission spectra were recorder in low volume 384-well black plates with clear bottom (Corning) for pFTAA (465nm – 600nm) and curcumin (450nm – 650nm), after exciting at 440nm and 420nm, respectively, using a ClarioStar plate reader at 30oC (BMG Labtech, Germany). The acquired spectra were background subtracted and normalised. The derived normalised spectra were then subjected to principal component analysis using the prcomp function in R.

### Fourier-Transform infrared spectroscopy (FTIR)

Similar equimolar mixtures, variant-only and PHF6 APR-only preparations were used for FTIR measurements. Each sample was split into equal aliquots and allowed to incubate for 7 days at ambient conditions. Droplets (5μL) of peptide samples (n=4) were cast onto a 96-well silicon microplate (Bruker) and dried to form thin films. FTIR spectra were recorded as averages of 64 spectral scans at 4 nm^-1^ resolution in transmission mode to reduce signal-to-noise ratio, using an HTS-XT FTIR microplate reader (Bruker). Background correction was performed by subtracting spectra obtained from a blank position of the microplate. Spectral normalisation and 2^nd^ derivatives with a 13-point smoothing, using Savitzky-Golay filtering^128^, were calculated using the OPUS software after isolation of the amide I and amide II regions of the derived spectra (1700-1500cm^-1^). The normalised spectra were subjected to principal component analysis using the prcomp function in R.

### Fluorescence recovery after photobleaching (FRAP)

Confocal microscopy was used to acquire images for fluorescence recovery. Instances were acquired as individual frames on a Nikon A1R Eclipse Ti confocal microscope, equipped with a Plan APO VC 60x oil lens. For bleaching, we defined a region-of-interest (ROI) that was excited using the CFP-donor laser line (405nm) at 100% laser power and emission was collected using the YFP acceptor filter (550nm). FRAP was performed in pulses of successive time increments (0.06s, 0.6s and 1.2s). Between pulses, total fluorescence of the ROI was measured for 10s (ROI within spot) or 20s (ROI in the cellular background) by acquiring single frames every 2s (5 or 10 frames per window, respectively). Total fluorescence of the ROI in each frame was normalized to the total fluorescence of the same pre-bleached region (t=0) to check for potential recovery.

### Tau aggregation and seed preparation

Recombinant full-length tau (tau^2N4R^) was produced following previous established protocols^129^. Lyophilised protein aliquots were freshly dissolved in 10mM HEPES pH 7.4 supplemented with 100mM NaCl at a final concentration of 10μM. After filtration, using 0.2 μM PVDF filters, the protein solution was spiked with 5μM of heparin (Sigma) and aggregation was monitored by adding 25μM of Th-T in half-area black 96-well microplates (Corning, USA). Fluorescence was measured in triplicates, using a FluoStar Omega plate reader (BMG Labtech, Germany) at 37°C, equipped with an excitation filter at 440nm and emission filter at 490nm. To generate seeds, endpoint amyloid fibrils were sonicated for 15min (30s on, 30s off) at 10°C, using a Bioruptor Pico sonication device (Diagenode).

### FRET cellular transfection assays

HEK293 Tau^RD^-P301S-CFP/YFP expressing biosensor cells^98^ were purchased form ATCC and cultured in DMEM medium, supplemented with 10% FBS at 37°C, and a 5% CO2 atmosphere. Gene constructs (**Table 2**) were generated and onboarded to a pTwist CMV expression vector by coupling double-tagged (N-terminal HA and C-terminal FLAG recognition sites) genes of interest to an mKO2 fluorescence reporter, separated by an IRES site to enable independent co-expression (Twist Biosciences). Due to restrictions imposed by the construct synthesis (Twist Biosciences), for proteins longer than 500 residues we designed shorter domain-constructs containing the homologous sequences (**Table 2**). Biosensor cells were plated in poly-L-lysine coated 96-well plates (PerkinElmer) at a density of 20000 cells/well. DNA transfection (100ng) and tau seeds transfection was performed 6h and 48h later, using Lipofectamine 3000 according to the manufacturer guidelines and cells were fixed with 4% formaldehyde 24h after seeding. Fixed cells were stained with DAPI (Thermofisher, D1306) following the manufacturer protocol. For immunofluorescence staining, primary antibody staining at 1:1000 dilution was performed in 1% BSA with an HA-tag (C29F4) Rabbit mAb (Cell Signalling, #3724), followed by secondary staining with an Alexa Fluor 647 goat anti-rabbit antibody (ThermoFisher, A-21245) at 1:1000 dilution in 1% BSA for 1h. Three individual plate preparations were performed for each construct gradient as independent experiments (n=3). High-content screening was performed at the VIB Screening Core/C-BIOS, using an Opera Phenix HCS (PerkinElmer) equipped with proper filter channels to track tau aggregation through FRET (Ex:405, Em:550), construct colocalization through HA-staining (Ex:647, Em:667) and DAPI staining (Ex:405 Em:430). Image storage and segmentation analysis was performed using the Columbus Plus digital platform (PerkinElmer).

### Microscale Thermophoresis (MST)

MST measurements were performed to calculate binding affinities. Monomeric tau was labelled using amine reactive ATTO_633_ (ATTO_633_-NHS), following the manufacturer guidelines. Labelled tau aggregates were prepared using a 1:9 analogy of labelled to unlabelled monomeric tau, following the same aggregation protocol described for the unlabelled protein. 25nM of ATTO_633_-monomeric tau or ATTO_633_-tau seeds were mixed against the CAP1 inhibitor, which was dissolved and titrated down starting from 50μM, in tau buffer (HEPES 10mM, 100mM NaCl). Measurements were recorded on a Monolith NT automated instrument (NanoTemper Technologies GmbH, Germany) with a red-laser channel at 5% LED excitation power and medium MST power at ambient conditions. Affinity constants and experimental data fitting was performed using the NanoTemper analysis software (v2.2.4) and results were depicted as differentials between the bound and unbound state after baseline subtraction (ΔFnorm) over inhibitor concentration in the logarithmic scale.

### Structure-based inhibition using FRET tau biosensor cells

We co-incubated tau seeds, at a concentration of 125nM, produced from recombinant full-length tau^2N4R^ as described above, with a titrated concentration gradient of the CAP1 peptide for 2h at room temperature. HEK293 Tau^RD^-P301S-CFP/YFP expressing biosensor cells were plated in poly-L-lysine coated 96-well plates (PerkinElmer) at a density of 20000 cells/well and subsequently transfected with pre-incubated mixtures of tau seeds/CAP1, using Lipofectamine 3000 according to the manufacturer guidelines. Cell fixation was performed 24h after transfection using 4% formaldehyde and cellular imaging was performed using an Operetta CLS (PerkinElmer). Three individual plate preparations were used as independent experiments for statistical significance (n=3). Data storage and analysis was performed using the Columbus Plus digital platform (PerkinElmer).

## Acknowledgements

We are grateful to the Diamond lab (University of Texas Southwestern Medical Center, Dallas, Texas, USA) for sharing their tau expression vector. The authors gratefully acknowledge Electron Microscopy Platform & Bio Imaging Core, Department of Neurosciences KU Leuven, VIB – KU Leuven Center for Brain & Disease Research for their support & assistance in this work. Nikon A1R Eclipse Ti confocal was acquired through a Hercules type 1 AKUL/097/037 grant to Wim Annaert. We also thank the VIB Screening Core/C-BIOS facility for excellent support with our cellular screens. This work was supported by the Flanders institute for Biotechnology (VIB); KU Leuven; the Fund for Scientific Research Flanders (FWO, project grants G0C2818N, G0C0320N, and G053420N, FWO/Hercules Foundation equipment grants FWO AKUL/15/34 -G0H1716N, I011620N, and NextGenQBio -AH2016.133, and Postdoctoral Fellowships 12P0919N and 12P0922N to NL); the Stichting Alzheimer Onderzoek (SAO-FRA 2019/0015, SAO-FRA 2020/0009, and SAO-FRA 2020/0013); and a VLAIO Innovation Mandate to EM (HBC.2020.2854).

## Author contributions

N.L., M.R, E.M., K.K., C.M., T.G. performed the *in vitro* experiments. N.L. performed the in silico analysis. N.L. and M.R. synthesized and purified the peptides. N.L., M.R., E.M., S.D’H., V.G. and D.A performed the in-cell experiments. N.L., E.M., V.G., D.A., F.R. and J.S. participated in analysing the data. N.L., F.R. and J.S. conceived and designed the study and prepared the manuscript. All authors approved the manuscript.

## Conflicts of interest

The authors declare no conflicts of interest

## Abbreviations

APRs: aggregation prone regions
PCA: principal component analysis
UMAP: uniform manifold approximation and projection
AD: alzheimer’s disease
PD: parkinson’s Disease
Aβ: beta-amyloid peptide
Apo-AI: apolipoprotein A-I
HA-tag: human influenza hemagglutinin tag
IRES: internal ribosome entry site
PHF: pair helical fragment
CFP: cyan-fluorescent protein
YFP: yellow-fluorescent protein
FRET: Förster resonance energy transfer
CBD: corticobasal degeneration
CJD: Creutzfeldt-Jakob disease
FTD: frontotemporal dementia
TEM: transmission Electron Microscopy
FTIR: Fourier-transform infrared spectroscopy
cryo-EM: cryogenic electron microscopy
NMR: nuclear magnetic resonance
CMV: cytomegalovirus
PVDF: polyvinylidene fluoride
FRAP: fluorescence recovery after photobleaching
T2D: type-2 diabetes
RP-HPLC: reverse-phase high-performance liquid chromatography
BSA: bovine serum albumin
NHS: succinimidyl ester
DOCK3: dedicator of cytokinesis 3
IDE: insulin-degrading enzyme
TRA2B: transformer-2 protein homolog beta
SNTG1: synaptotagmin-1
Hsp70: heat-shock protein 70
iPSCs: induced pluripotent stem cells

## Notes

### Competing Interest Statement

The authors have declared no competing interest.

### Summary of Updates

Author names corrected; Acknowledgements updated

## References

1 Van Dam, D., Vermeiren, Y., Dekker, A. D., Naude, P. J. & Deyn, P. P. Neuropsychiatric Disturbances in Alzheimer’s Disease: What Have We Learned from Neuropathological Studies? Curr Alzheimer Res 13, 1145–1164 (2016).

2 Dugger, B. N. & Dickson, D. W. Pathology of Neurodegenerative Diseases. Cold Spring Harbor perspectives in biology 9, doi:10.1101/cshperspect.a028035 (2017).

3 Eisenberg, D. & Jucker, M. The amyloid state of proteins in human diseases. Cell 148, 1188–1203, doi:10.1016/j.cell.2012.02.022 (2012).

4 Chiti, F. & Dobson, C. M. Protein Misfolding, Amyloid Formation, and Human Disease: A Summary of Progress Over the Last Decade. Annu Rev Biochem 86, 27–68, doi:10.1146/annurev-biochem-061516-045115 (2017).

5 Benson, M. D. et al. Amyloid nomenclature 2018: recommendations by the International Society of Amyloidosis (ISA) nomenclature committee. Amyloid 25, 215–219, doi:10.1080/13506129.2018.1549825 (2018).

6 Braak, H. et al. Amyotrophic lateral sclerosis--a model of corticofugal axonal spread. Nature reviews. Neurology 9, 708–714, doi:10.1038/nrneurol.2013.221 (2013).

7 Goedert, M., Eisenberg, D. S. & Crowther, R. A. Propagation of Tau Aggregates and Neurodegeneration. Annu Rev Neurosci 40, 189–210, doi:10.1146/annurev-neuro-072116-031153 (2017).

8 Braak, H., Alafuzoff, I., Arzberger, T., Kretzschmar, H. & Del Tredici, K. Staging of Alzheimer disease-associated neurofibrillary pathology using paraffin sections and immunocytochemistry. Acta Neuropathol 112, 389–404, doi:10.1007/s00401-006-0127-z (2006).

9 Fu, H., Hardy, J. & Duff, K. E. Selective vulnerability in neurodegenerative diseases. Nat Neurosci 21, 1350–1358, doi:10.1038/s41593-018-0221-2 (2018).

10 Bussiere, T. et al. Progressive degeneration of nonphosphorylated neurofilament protein-enriched pyramidal neurons predicts cognitive impairment in Alzheimer’s disease: stereologic analysis of prefrontal cortex area 9. J Comp Neurol 463, 281–302, doi:10.1002/cne.10760 (2003).

11 Sepulcre, J. et al. Hierarchical Organization of Tau and Amyloid Deposits in the Cerebral Cortex. JAMA Neurol 74, 813–820, doi:10.1001/jamaneurol.2017.0263 (2017).

12 Hardy, J. Catastrophic cliffs: a partial suggestion for selective vulnerability in neurodegenerative diseases. Biochem Soc Trans 44, 659–661, doi:10.1042/BST20150287 (2016).

13 Roussarie, J. P. et al. Selective Neuronal Vulnerability in Alzheimer’s Disease: A Network-Based Analysis. Neuron 107, 821–835 e812, doi:10.1016/j.neuron.2020.06.010 (2020).

14 Muratore, C. R. et al. Cell-type Dependent Alzheimer’s Disease Phenotypes: Probing the Biology of Selective Neuronal Vulnerability. Stem Cell Reports 9, 1868–1884, doi:10.1016/j.stemcr.2017.10.015 (2017).

15 Jaunmuktane, Z. & Brandner, S. On the journey to uncover the causes of selective cellular and regional vulnerability in neurodegeneration. Acta Neuropathol 138, 677–680, doi:10.1007/s00401-019-02079-9 (2019).

16 Akila Parvathy Dharshini, S., Taguchi, Y. H. & Michael Gromiha, M. Exploring the selective vulnerability in Alzheimer disease using tissue specific variant analysis. Genomics 111, 936–949, doi:10.1016/j.ygeno.2018.05.024 (2019).

17 Keo, A. et al. Transcriptomic signatures of brain regional vulnerability to Parkinson’s disease. Commun Biol 3, 101, doi:10.1038/s42003-020-0804-9 (2020).

18 Saxena, S. & Caroni, P. Selective neuronal vulnerability in neurodegenerative diseases: from stressor thresholds to degeneration. Neuron 71, 35–48, doi:10.1016/j.neuron.2011.06.031 (2011).

19 Konstantoulea, K., Louros, N., Rousseau, F. & Schymkowitz, J. Heterotypic interactions in amyloid function and disease. FEBS J, doi:10.1111/febs.15719 (2021).

20 Chin, J. et al. Reelin depletion in the entorhinal cortex of human amyloid precursor protein transgenic mice and humans with Alzheimer’s disease. J Neurosci 27, 2727–2733, doi:10.1523/JNEUROSCI.3758-06.2007 (2007).

21 Pujadas, L. et al. Reelin delays amyloid-beta fibril formation and rescues cognitive deficits in a model of Alzheimer’s disease. Nature Communications 5, doi:ARTN 3443 10.1038/ncomms4443 (2014).

22 Cuchillo-Ibanez, I. et al. The beta-amyloid peptide compromises Reelin signaling in Alzheimer’s disease. Sci Rep 6, 31646, doi:10.1038/srep31646 (2016).

23 Banerjee, S., Ferdosh, S., Ghosh, A. N. & Barat, C. Tau protein-induced sequestration of the eukaryotic ribosome: Implications in neurodegenerative disease. Sci Rep 10, 5225, doi:10.1038/s41598-020-61777-7 (2020).

24 Pathak, B. K., Mondal, S., Banerjee, S., Ghosh, A. N. & Barat, C. Sequestration of Ribosome during Protein Aggregate Formation: Contribution of ribosomal RNA. Sci Rep 7, 42017, doi:10.1038/srep42017 (2017).

25 Azizi, S. A. & Azizi, S. A. Synucleinopathies in neurodegenerative diseases: Accomplices, an inside job and selective vulnerability. Neurosci Lett 672, 150–152, doi:10.1016/j.neulet.2017.12.003 (2018).

26 Horvath, I. et al. Co-aggregation of pro-inflammatory S100A9 with alpha-synuclein in Parkinson’s disease: ex vivo and in vitro studies. J Neuroinflammation 15, 172, doi:10.1186/s12974-018-1210-9 (2018).

27 Ryan, B. J., Hoek, S., Fon, E. A. & Wade-Martins, R. Mitochondrial dysfunction and mitophagy in Parkinson’s: from familial to sporadic disease. Trends Biochem Sci 40, 200–210, doi:10.1016/j.tibs.2015.02.003 (2015).

28 Ciryam, P., Kundra, R., Morimoto, R. I., Dobson, C. M. & Vendruscolo, M. Supersaturation is a major driving force for protein aggregation in neurodegenerative diseases. Trends Pharmacol Sci 36, 72–77, doi:10.1016/j.tips.2014.12.004 (2015).

29 Hardy, J. Expression of normal sequence pathogenic proteins for neurodegenerative disease contributes to disease risk: ‘permissive templating’ as a general mechanism underlying neurodegeneration. Biochem Soc Trans 33, 578–581, doi:10.1042/BST0330578 (2005).

30 Ciryam, P., Tartaglia, G. G., Morimoto, R. I., Dobson, C. M. & Vendruscolo, M. Widespread aggregation and neurodegenerative diseases are associated with supersaturated proteins. Cell reports 5, 781–790, doi:10.1016/j.celrep.2013.09.043 (2013).

31 Freer, R. et al. Supersaturated proteins are enriched at synapses and underlie cell and tissue vulnerability in Alzheimer’s disease. Heliyon 5, e02589, doi:10.1016/j.heliyon.2019.e02589 (2019).

32 Ciryam, P. et al. A transcriptional signature of Alzheimer’s disease is associated with a metastable subproteome at risk for aggregation. Proc Natl Acad Sci U S A 113, 4753–4758, doi:10.1073/pnas.1516604113 (2016).

33 Fitzpatrick, A. W. P. et al. Cryo-EM structures of tau filaments from Alzheimer’s disease. Nature 547, 185–190, doi:10.1038/nature23002 (2017).

34 Scheres, S. H., Zhang, W., Falcon, B. & Goedert, M. Cryo-EM structures of tau filaments. Curr Opin Struct Biol 64, 17–25, doi:10.1016/j.sbi.2020.05.011 (2020).

35 Strohaker, T. et al. Structural heterogeneity of alpha-synuclein fibrils amplified from patient brain extracts. Nat Commun 10, 5535, doi:10.1038/s41467-019-13564-w (2019).

36 Landreh, M. et al. The formation, function and regulation of amyloids: insights from structural biology. J Intern Med 280, 164–176, doi:10.1111/joim.12500 (2016).

37 Riek, R. & Eisenberg, D. S. The activities of amyloids from a structural perspective. Nature 539, 227–235, doi:10.1038/nature20416 (2016).

38 Lutter, L., Serpell, C. J., Tuite, M. F. & Xue, W. F. The molecular lifecycle of amyloid -Mechanism of assembly, mesoscopic organisation, polymorphism, suprastructures, and biological consequences. Biochim Biophys Acta Proteins Proteom 1867, 140257, doi:10.1016/j.bbapap.2019.07.010 (2019).

39 Gallardo, R., Ranson, N. A. & Radford, S. E. Amyloid structures: much more than just a cross-beta fold. Curr Opin Struct Biol 60, 7–16, doi:10.1016/j.sbi.2019.09.001 (2020).

40 van der Kant, R., Louros, N., Schymkowitz, J. & Rousseau, F. A structural analysis of amyloid polymorphism in disease: clues for selective vulnerability? bioRxiv, 2021.2003.2001.433317, doi:10.1101/2021.03.01.433317 (2021).

41 Marshall, K. E. et al. A critical role for the self-assembly of Amyloid-beta1-42 in neurodegeneration. Sci Rep 6, 30182, doi:10.1038/srep30182 (2016).

42 Ganesan, A. et al. Structural hot spots for the solubility of globular proteins. Nature communications 7, 10816, doi:10.1038/ncomms10816 (2016).

43 Ventura, S. et al. Short amino acid stretches can mediate amyloid formation in globular proteins: the Src homology 3 (SH3) case. Proc Natl Acad Sci U S A 101, 7258–7263, doi:10.1073/pnas.03082491010308249101[pii] (2004).

44 Teng, P. K. & Eisenberg, D. Short protein segments can drive a non-fibrillizing protein into the amyloid state. Protein Engineering Design & Selection 22, 531–536, doi:10.1093/protein/gzp037 (2009).

45 Louros, N., Orlando, G., De Vleeschouwer, M., Rousseau, F. & Schymkowitz, J. Structure-based machine-guided mapping of amyloid sequence space reveals uncharted sequence clusters with higher solubilities. Nat Commun 11, 3314, doi:10.1038/s41467-020-17207-3 (2020).

46 Wetzel, R. Kinetics and thermodynamics of amyloid fibril assembly. Acc Chem Res 39, 671–679, doi:10.1021/ar050069h (2006).

47 O’Nuallain, B., Shivaprasad, S., Kheterpal, I. & Wetzel, R. Thermodynamics of A beta(1-40) amyloid fibril elongation. Biochemistry 44, 12709–12718 (2005).

48 O’Nuallain, B., Williams, A. D., Westermark, P. & Wetzel, R. Seeding specificity in amyloid growth induced by heterologous fibrils. J Biol Chem 279, 17490–17499 (2004).

49 Krebs, M. R., Morozova-Roche, L. A., Daniel, K., Robinson, C. V. & Dobson, C. M. Observation of sequence specificity in the seeding of protein amyloid fibrils. Protein Sci 13, 1933–1938, doi:10.1110/ps.0470700413/7/1933[pii] (2004).

50 Vanik, D. L., Surewicz, K. A. & Surewicz, W. K. Molecular basis of barriers for interspecies transmissibility of mammalian prions. Mol Cell 14, 139–145, doi:S1097276504001558 [pii] (2004).

51 Giasson, B. I. et al. Initiation and Synergistic Fibrillization of Tau and Alpha-Synuclein. Science 300, 636–640, doi:10.1126/science.1082324 (2003).

52 Oskarsson, M. E. et al. In vivo seeding and cross-seeding of localized amyloidosis: a molecular link between type 2 diabetes and Alzheimer disease. Am J Pathol 185, 834–846, doi:10.1016/j.ajpath.2014.11.016 (2015).

53 Ly, H. et al. The association of circulating amylin with β-amyloid in familial Alzheimer’s disease. Alzheimer’s & Dementia: Translational Research & Clinical Interventions 7, e12130, doi:https://doi.org/10.1002/trc2.12130 (2021).

54 Apostol, M. I., Wiltzius, J. J., Sawaya, M. R., Cascio, D. & Eisenberg, D. Atomic structures suggest determinants of transmission barriers in mammalian prion disease. Biochemistry 50, 2456–2463, doi:10.1021/bi101803k (2011).

55 Krotee, P. et al. Common fibrillar spines of amyloid-beta and human islet amyloid polypeptide revealed by microelectron diffraction and structure-based inhibitors. J Biol Chem 293, 2888–2902, doi:10.1074/jbc.M117.806109 (2018).

56 Vasconcelos, B. et al. Heterotypic seeding of Tau fibrillization by pre-aggregated Abeta provides potent seeds for prion-like seeding and propagation of Tau-pathology in vivo. Acta Neuropathol 131, 549–569, doi:10.1007/s00401-015-1525-x (2016).

57 Colom-Cadena, M. et al. Confluence of alpha-synuclein, tau, and beta-amyloid pathologies in dementia with Lewy bodies. J Neuropathol Exp Neurol 72, 1203–1212, doi:10.1097/NEN.0000000000000018 (2013).

58 Pham, C. L. et al. Viral M45 and necroptosis-associated proteins form heteromeric amyloid assemblies. EMBO Rep 20, doi:10.15252/embr.201846518 (2019).

59 Sampson, T. R. et al. A gut bacterial amyloid promotes alpha-synuclein aggregation and motor impairment in mice. Elife 9, doi:10.7554/eLife.53111 (2020).

60 Collinson, S. K., Parker, J. M., Hodges, R. S. & Kay, W. W. Structural predictions of AgfA, the insoluble fimbrial subunit of Salmonella thin aggregative fimbriae. Journal of molecular biology 290, 741–756, doi:10.1006/jmbi.1999.2882 (1999).

61 Louros, N. N., Bolas, G. M. P., Tsiolaki, P. L., Hamodrakas, S. J. & Iconomidou, V. A. Intrinsic aggregation propensity of the CsgB nucleator protein is crucial for curli fiber formation. J Struct Biol 195, 179–189, doi:10.1016/j.jsb.2016.05.012 (2016).

62 White, A. P. et al. Structure and characterization of AgfB from Salmonella enteritidis thin aggregative fimbriae. Journal of molecular biology 311, 735–749, doi:10.1006/jmbi.2001.4876 (2001).

63 Louros, N. N. et al. A common ‘aggregation-prone’ interface possibly participates in the self-assembly of human zona pellucida proteins. FEBS Lett 590, 619–630, doi:10.1002/1873-3468.12099 (2016).

64 Li, J. et al. The RIP1/RIP3 necrosome forms a functional amyloid signaling complex required for programmed necrosis. Cell 150, 339–350, doi:10.1016/j.cell.2012.06.019 (2012).

65 Wu, X. N. et al. Distinct roles of RIP1-RIP3 hetero- and RIP3-RIP3 homo-interaction in mediating necroptosis. Cell Death Differ 21, 1709–1720, doi:10.1038/cdd.2014.77 (2014).

66 Petrie, E. J. et al. Viral MLKL Homologs Subvert Necroptotic Cell Death by Sequestering Cellular RIPK3. Cell Rep 28, 3309–3319 e3305, doi:10.1016/j.celrep.2019.08.055 (2019).

67 Wolozin, B. Regulated protein aggregation: stress granules and neurodegeneration. Mol Neurodegener 7, 56, doi:10.1186/1750-1326-7-56 (2012).

68 Wolozin, B. & Ivanov, P. Stress granules and neurodegeneration. Nat Rev Neurosci 20, 649–666, doi:10.1038/s41583-019-0222-5 (2019).

69 Fabiani, C. & Antollini, S. S. Alzheimer’s Disease as a Membrane Disorder: Spatial Cross-Talk Among Beta-Amyloid Peptides, Nicotinic Acetylcholine Receptors and Lipid Rafts. Front Cell Neurosci 13, 309, doi:10.3389/fncel.2019.00309 (2019).

70 Stewart, K. L. et al. Atomic Details of the Interactions of Glycosaminoglycans with Amyloid-beta Fibrils. J Am Chem Soc 138, 8328–8331, doi:10.1021/jacs.6b02816 (2016).

71 Cohen, S. I. A. et al. A molecular chaperone breaks the catalytic cycle that generates toxic Abeta oligomers. Nat Struct Mol Biol 22, 207–213, doi:10.1038/nsmb.2971 (2015).

72 Derkatch, I. L. et al. Effects of Q/N-rich, polyQ, and non-polyQ amyloids on the de novo formation of the [PSI+] prion in yeast and aggregation of Sup35 in vitro. Proc Natl Acad Sci U S A 101, 12934–12939, doi:10.1073/pnas.0404968101 (2004).

73 Mompean, M. et al. The Structure of the Necrosome RIPK1-RIPK3 Core, a Human Hetero-Amyloid Signaling Complex. Cell 173, 1244–1253 e1210, doi:10.1016/j.cell.2018.03.032 (2018).

74 Sidhu, A., Segers-Nolten, I. & Subramaniam, V. Conformational Compatibility Is Essential for Heterologous Aggregation of alpha-Synuclein. ACS Chem Neurosci 7, 719–727, doi:10.1021/acschemneuro.5b00322 (2016).

75 Wasmer, C. et al. Structural similarity between the prion domain of HET-s and a homologue can explain amyloid cross-seeding in spite of limited sequence identity. Journal of molecular biology 402, 311–325, doi:10.1016/j.jmb.2010.06.053 (2010).

76 Benkemoun, L. et al. Two structurally similar fungal prions efficiently cross-seed in vivo but form distinct polymers when coexpressed. Mol Microbiol 82, 1392–1405, doi:10.1111/j.1365-2958.2011.07893.x (2011).

77 Schymkowitz, J. et al. The FoldX web server: an online force field. Nucleic Acids Res 33, W382–388, doi:10.1093/nar/gki387 (2005).

78 Khodaparast, L. et al. Aggregating sequences that occur in many proteins constitute weak spots of bacterial proteostasis. Nat Commun 9, 866, doi:10.1038/s41467-018-03131-0 (2018).

79 Michiels, E. et al. Reverse engineering synthetic antiviral amyloids. Nat Commun 11, 2832, doi:10.1038/s41467-020-16721-8 (2020).

80 Houben, B. et al. Autonomous aggregation suppression by acidic residues explains why chaperones favour basic residues. EMBO J 39, e102864, doi:10.15252/embj.2019102864 (2020).

81 Richardson, J. S. & Richardson, D. C. Natural beta-sheet proteins use negative design to avoid edge-to-edge aggregation. Proc Natl Acad Sci U S A 99, 2754–2759, doi:10.1073/pnas.052706099 (2002).

82 Banach, M. & Roterman, I. in From Globular Proteins to Amyloids (ed Irena Roterman-Konieczna) 95–115 (Elsevier, 2020).

83 Kajava, A. V. & Steven, A. C. Beta-rolls, beta-helices, and other beta-solenoid proteins. Adv Protein Chem 73, 55–96, doi:10.1016/S0065-3233(06)73003-0 (2006).

84 Louros, N. N., Baltoumas, F. A., Hamodrakas, S. J. & Iconomidou, V. A. A beta-solenoid model of the Pmel17 repeat domain: insights to the formation of functional amyloid fibrils. J Comput Aided Mol Des 30, 153–164, doi:10.1007/s10822-015-9892-x (2016).

85 Sawaya, M. R. et al. Atomic structures of amyloid cross-beta spines reveal varied steric zippers. Nature 447, 453–457, doi:10.1038/nature05695 (2007).

86 von Bergen, M., Barghorn, S., Biernat, J., Mandelkow, E. M. & Mandelkow, E. Tau aggregation is driven by a transition from random coil to beta sheet structure. Biochim Biophys Acta 1739, 158–166, doi:10.1016/j.bbadis.2004.09.010 (2005).

87 Seidler, P. M. et al. Structure-based inhibitors of tau aggregation. Nat Chem 10, 170–176, doi:10.1038/nchem.2889 (2018).

88 Louros, N. et al. WALTZ-DB 2.0: an updated database containing structural information of experimentally determined amyloid-forming peptides. Nucleic Acids Res 48, D389–D393, doi:10.1093/nar/gkz758 (2020).

89 Louros, N. N. et al. Chameleon ‘aggregation-prone’ segments of apoA-I: A model of amyloid fibrils formed in apoA-I amyloidosis. Int J Biol Macromol 79, 711–718, doi:10.1016/j.ijbiomac.2015.05.032 (2015).

90 Blancas-Mejia, L. M. & Ramirez-Alvarado, M. Recruitment of Light Chains by Homologous and Heterologous Fibrils Shows Distinctive Kinetic and Conformational Specificity. Biochemistry 55, 2967–2978, doi:10.1021/acs.biochem.6b00090 (2016).

91 Liberta, F. et al. Cryo-EM fibril structures from systemic AA amyloidosis reveal the species complementarity of pathological amyloids. Nat Commun 10, 1104, doi:10.1038/s41467-019-09033-z (2019).

92 Boyer, D. R. et al. The alpha-synuclein hereditary mutation E46K unlocks a more stable, pathogenic fibril structure. Proc Natl Acad Sci U S A 117, 3592–3602, doi:10.1073/pnas.1917914117 (2020).

93 Sun, Y. et al. Cryo-EM structure of full-length alpha-synuclein amyloid fibril with Parkinson’s disease familial A53T mutation. Cell Res 30, 360–362, doi:10.1038/s41422-020-0299-4 (2020).

94 Sgourakis, N. G., Yau, W. M. & Qiang, W. Modeling an in-register, parallel “iowa” abeta fibril structure using solid-state NMR data from labeled samples with rosetta. Structure 23, 216–227, doi:10.1016/j.str.2014.10.022 (2015).

95 Schutz, A. K. et al. Atomic-resolution three-dimensional structure of amyloid beta fibrils bearing the Osaka mutation. Angew Chem Int Ed Engl 54, 331–335, doi:10.1002/anie.201408598 (2015).

96 Gallardo, R. et al. Fibril structures of diabetes-related amylin variants reveal a basis for surface-templated assembly. Nature Structural & Molecular Biology, doi:10.1038/s41594-020-0496-3 (2020).

97 Condello, C. et al. Structural heterogeneity and intersubject variability of Abeta in familial and sporadic Alzheimer’s disease. Proc Natl Acad Sci U S A 115, E782–E791, doi:10.1073/pnas.1714966115 (2018).

98 Holmes, B. B. et al. Proteopathic tau seeding predicts tauopathy in vivo. Proc Natl Acad Sci U S A 111, E4376–4385, doi:10.1073/pnas.1411649111 (2014).

99 Kar, K., Arduini, I., Drombosky, K. W., van der Wel, P. C. & Wetzel, R. D-polyglutamine amyloid recruits L-polyglutamine monomers and kills cells. Journal of molecular biology 426, 816–829, doi:10.1016/j.jmb.2013.11.019 (2014).

100 Kapurniotu, A., Schmauder, A. & Tenidis, K. Structure-based design and study of non-amyloidogenic, double N-methylated IAPP amyloid core sequences as inhibitors of IAPP amyloid formation and cytotoxicity. Journal of molecular biology 315, 339–350, doi:10.1006/jmbi.2001.5244 (2002).

101 Tatarek-Nossol, M. et al. Inhibition of hIAPP amyloid-fibril formation and apoptotic cell death by a designed hIAPP amyloid-core-containing hexapeptide. Chem Biol 12, 797–809, doi:10.1016/j.chembiol.2005.05.010 (2005).

102 Yan, L. M. et al. Selectively N-methylated soluble IAPP mimics as potent IAPP receptor agonists and nanomolar inhibitors of cytotoxic self-assembly of both IAPP and Abeta40. Angew Chem Int Ed Engl 52, 10378–10383, doi:10.1002/anie.201302840 (2013).

103 Sangwan, S. et al. Inhibition of synucleinopathic seeding by rationally designed inhibitors. Elife 9, doi:10.7554/eLife.46775 (2020).

104 Seidler, P. M. et al. Structure-based inhibitors halt prion-like seeding by Alzheimer’s disease-and tauopathy-derived brain tissue samples. J Biol Chem 294, 16451–16464, doi:10.1074/jbc.RA119.009688 (2019).

105 Saelices, L. et al. A pair of peptides inhibits seeding of the hormone transporter transthyretin into amyloid fibrils. J Biol Chem 294, 6130–6141, doi:10.1074/jbc.RA118.005257 (2019).

106 Lu, J. et al. Structure-Based Peptide Inhibitor Design of Amyloid-beta Aggregation. Front Mol Neurosci 12, 54, doi:10.3389/fnmol.2019.00054 (2019).

107 Griner, S. L. et al. Structure-based inhibitors of amyloid beta core suggest a common interface with tau. Elife 8, doi:10.7554/eLife.46924 (2019).

108 Cheng, P. N., Liu, C., Zhao, M., Eisenberg, D. & Nowick, J. S. Amyloid beta-sheet mimics that antagonize protein aggregation and reduce amyloid toxicity. Nat Chem4, 927–933, doi:10.1038/nchem.1433 (2012).

109 Gallardo, R. et al. De novo design of a biologically active amyloid. Science 354, doi:10.1126/science.aah4949 (2016).

110 Mathieu, C., Pappu, R. V. & Taylor, J. P. Beyond aggregation: Pathological phase transitions in neurodegenerative disease. Science 370, 56–60, doi:10.1126/science.abb8032 (2020).

111 Brunello, C. A., Yan, X. & Huttunen, H. J. Internalized Tau sensitizes cells to stress by promoting formation and stability of stress granules. Sci Rep 6, 30498, doi:10.1038/srep30498 (2016).

112 Gui, X. et al. Structural basis for reversible amyloids of hnRNPA1 elucidates their role in stress granule assembly. Nat Commun 10, 2006, doi:10.1038/s41467-019-09902-7 (2019).

113 Ray, S. et al. alpha-Synuclein aggregation nucleates through liquid-liquid phase separation. Nat Chem 12, 705–716, doi:10.1038/s41557-020-0465-9 (2020).

114 Jha, N. K. et al. Impact of Insulin Degrading Enzyme and Neprilysin in Alzheimer’s Disease Biology: Characterization of Putative Cognates for Therapeutic Applications. J Alzheimers Dis 48, 891–917, doi:10.3233/JAD-150379 (2015).

115 Kurochkin, I. V., Guarnera, E. & Berezovsky, I. N. Insulin-Degrading Enzyme in the Fight against Alzheimer’s Disease. Trends Pharmacol Sci 39, 49–58, doi:10.1016/j.tips.2017.10.008 (2018).

116 Sahoo, B. R. et al. Degradation of Alzheimer’s Amyloid-beta by a Catalytically Inactive Insulin-Degrading Enzyme. J Mol Biol 433, 166993, doi:10.1016/j.jmb.2021.166993 (2021).

117 Chen, Q. et al. Presenilin binding protein is associated with neurofibrillary alterations in Alzheimer’s disease and stimulates tau phosphorylation. Am J Pathol 159, 1597–1602, doi:10.1016/S0002-9440(10)63005-2 (2001).

118 Liu, X. Y. et al. Regulation of RAGE splicing by hnRNP A1 and Tra2beta-1 and its potential role in AD pathogenesis. J Neurochem 133, 187–198, doi:10.1111/jnc.13069 (2015).

119 Wong, J. Altered expression of RNA splicing proteins in Alzheimer’s disease patients: evidence from two microarray studies. Dement Geriatr Cogn Dis Extra 3, 74–85, doi:10.1159/000348406 (2013).

120 Storbeck, M. et al. Neuronal-specific deficiency of the splicing factor Tra2b causes apoptosis in neurogenic areas of the developing mouse brain. PLoS One 9, e89020, doi:10.1371/journal.pone.0089020 (2014).

121 Lester, E. et al. Tau aggregates are RNA-protein assemblies that mislocalize multiple nuclear speckle components. Neuron, doi:10.1016/j.neuron.2021.03.026 (2021).

122 Öhrfelt, A. et al. The pre-synaptic vesicle protein synaptotagmin is a novel biomarker for Alzheimer’s disease. Alzheimer’s Research & Therapy 8, 41, doi:10.1186/s13195-016-0208-8 (2016).

123 Keller, L. J. et al. Presenilin 1 increases association with synaptotagmin 1 during normal aging. Neurobiology of Aging 86, 156–161, doi:https://doi.org/10.1016/j.neurobiolaging.2019.10.006 (2020).

124 Zoltowska, K. M. et al. Dynamic presenilin 1 and synaptotagmin 1 interaction modulates exocytosis and amyloid β production. Molecular Neurodegeneration 12, 15, doi:10.1186/s13024-017-0159-y (2017).

125 Iwasawa, S. et al. Recurrent de novo MAPK8IP3 variants cause neurological phenotypes. Ann Neurol 85, 927–933, doi:10.1002/ana.25481 (2019).

126 Gowrishankar, S. et al. Overlapping roles of JIP3 and JIP4 in promoting axonal transport of lysosomes in human iPSC-derived neurons. bioRxiv, 2020.2006.2013.149443, doi:10.1101/2020.06.13.149443 (2021).

127 Berman, H. M. et al. The Protein Data Bank. Nucleic acids research 28, 235–242, doi:10.1093/nar/28.1.235 (2000).

128 Savitzky, A. & Golay, M. J. E. Smoothing and Differentiation of Data by Simplified Least Squares Procedures. Analytical Chemistry 36, 1627–1639, doi:10.1021/ac60214a047 (1964).

129 Mirbaha, H. et al. Inert and seed-competent tau monomers suggest structural origins of aggregation. Elife 7, doi:10.7554/eLife.36584 (2018).

