## Supplementary information for "Mapping the sequence specificity of heterotypic amyloid interactions enables the identification of aggregation modifiers"

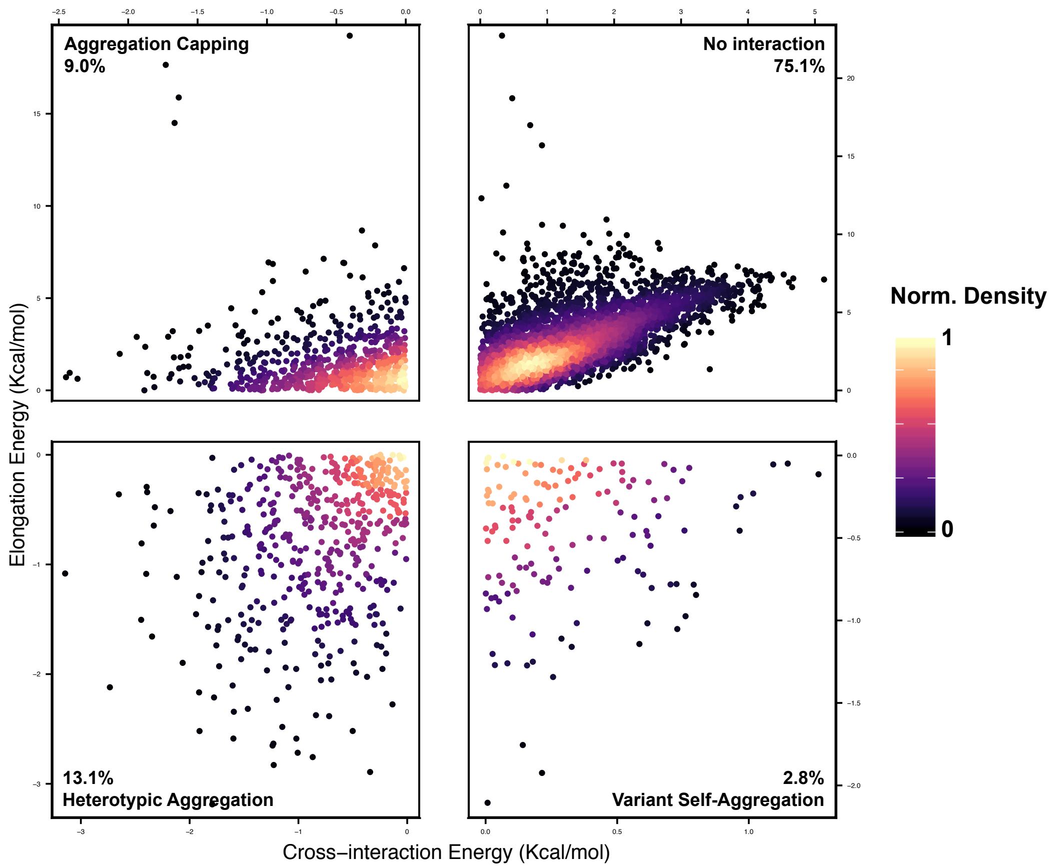


**Supplementary Figure 1. Thermodynamic profile of all possible double variants for VQIVYK.** Distribution analysis indicates that a smaller fraction of double variants is compatible to heterotypic or even self-associating interactions.

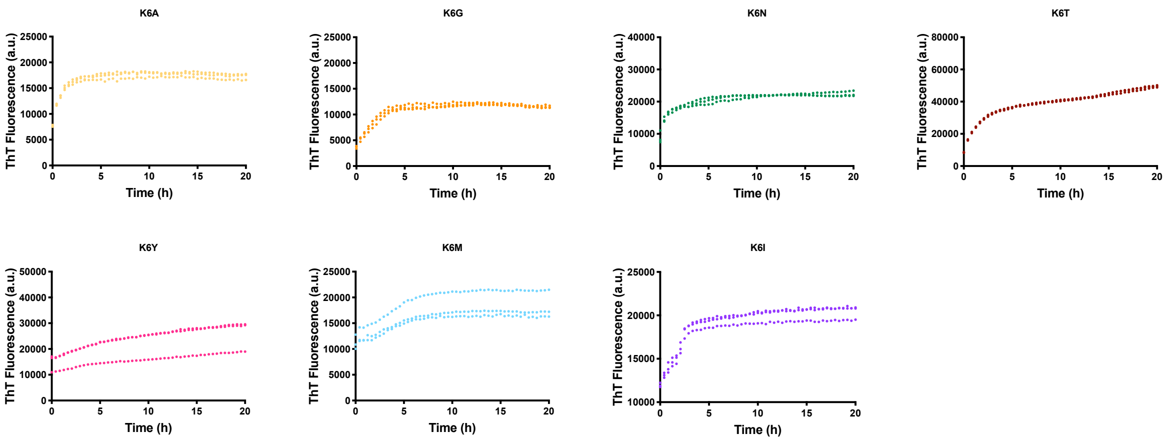


**Supplementary Figure 2.** Th-T kinetics, performed in triplicates at a concentration of 25μM, for variants corresponding to the exposed Lys residue of VQIVYK.

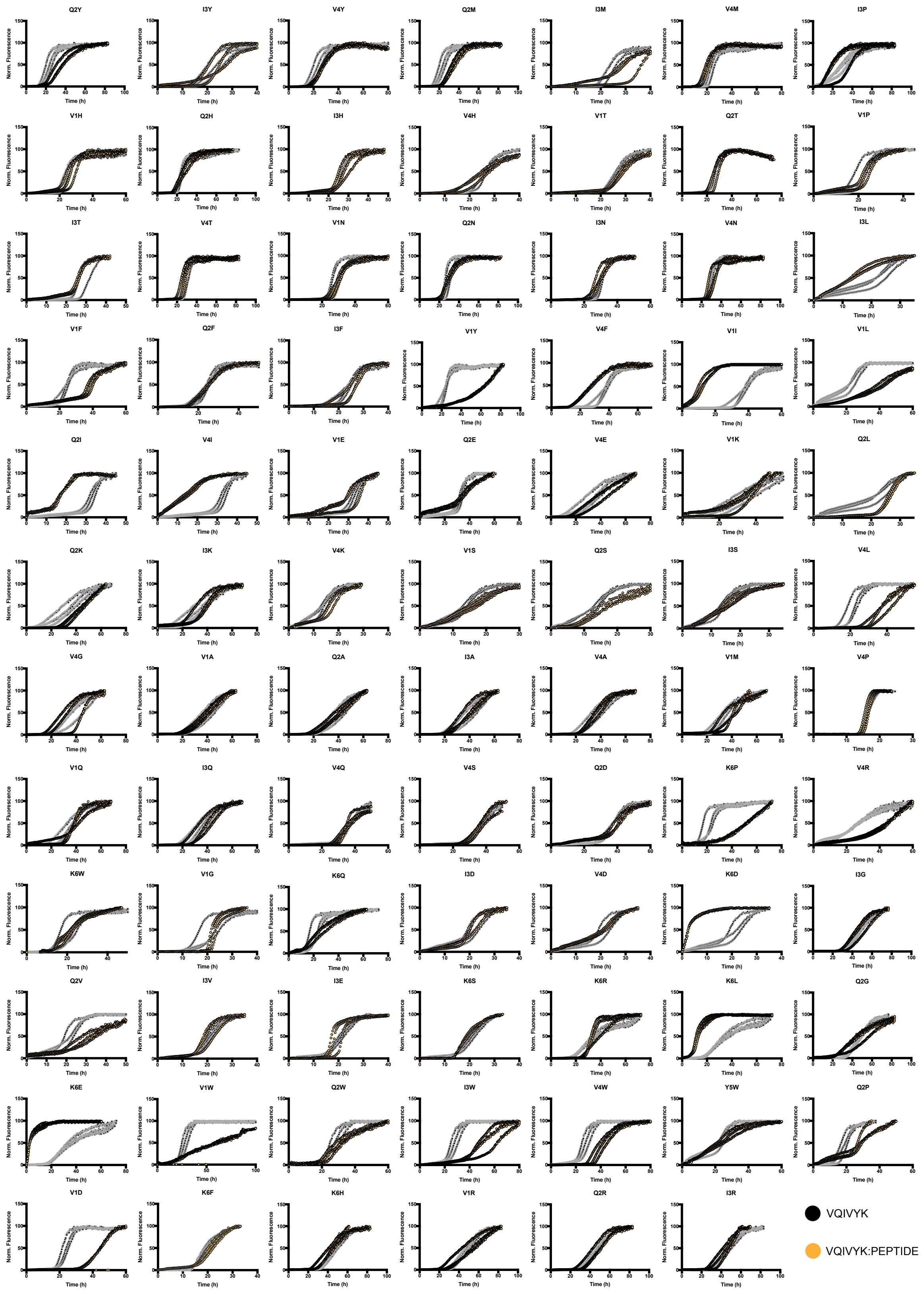


**Supplementary Figure 3.** Th-T curves of the VQIVYK peptide mixed (1:5) with the library variants.

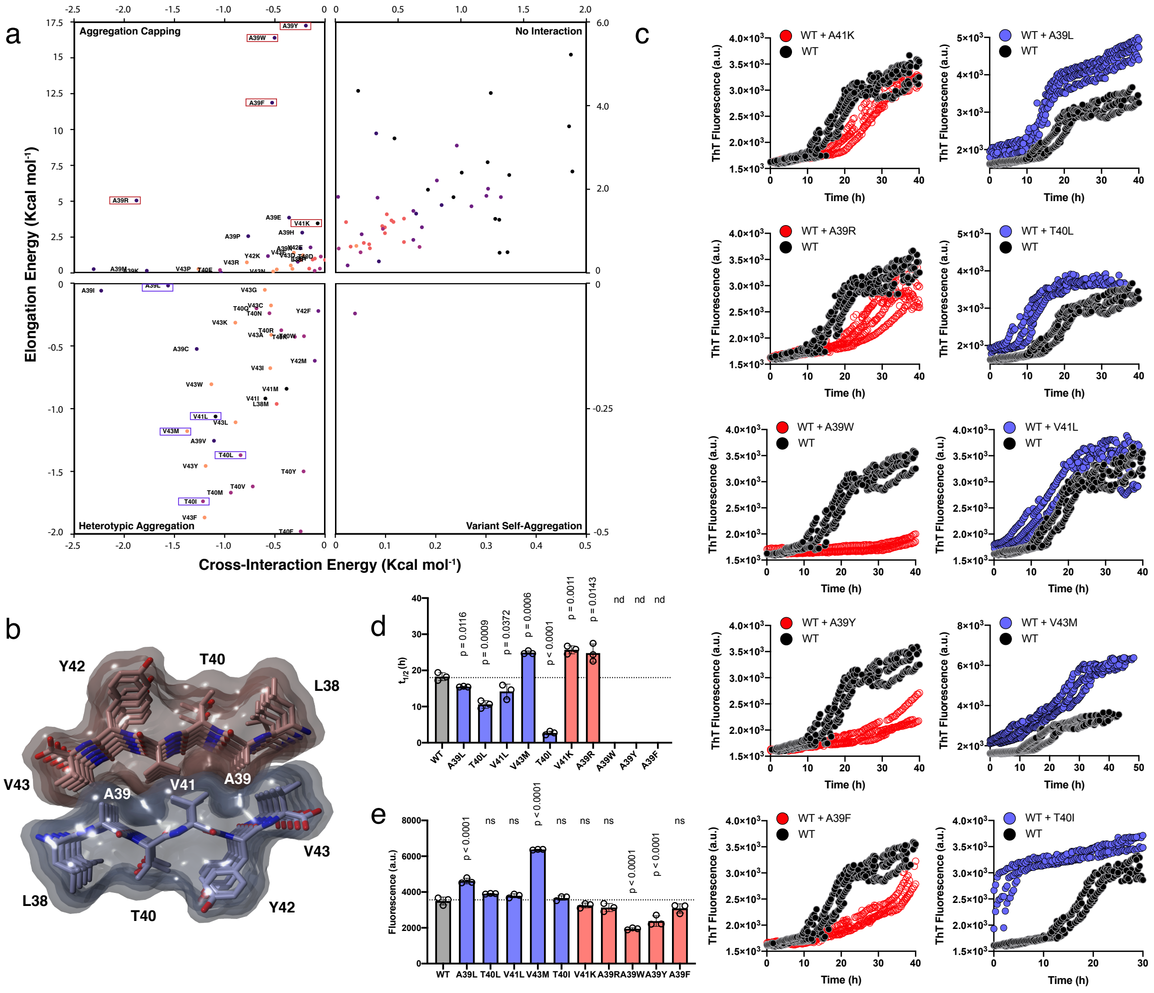

**Supplementary Figure 4. Thermodynamic profiling using a three-dimensional model template of the $^{38}$LATVYV$^{43}$ APR from ApoA-I.** (a) Thermodynamic profiling of cross-interaction propensities for single position homologs. Highlighted in red and purple boxes are randomly selected sequences that were experimentally tested. (b) Topological model of the aggregation core formed by the $^{38}$LATVYV$^{43}$ APR, developed using CORDAX[1]. (c) Th-T kinetics, performed in triplicates, as well as (d) halftime and (e) endpoint fluorescence analysis, validated that all 10 peptides either capped or promoted aggregation of the APR, as depicted by the thermodynamic analysis. Statistical significance was determined using Tukey's multiple comparisons to the WT.

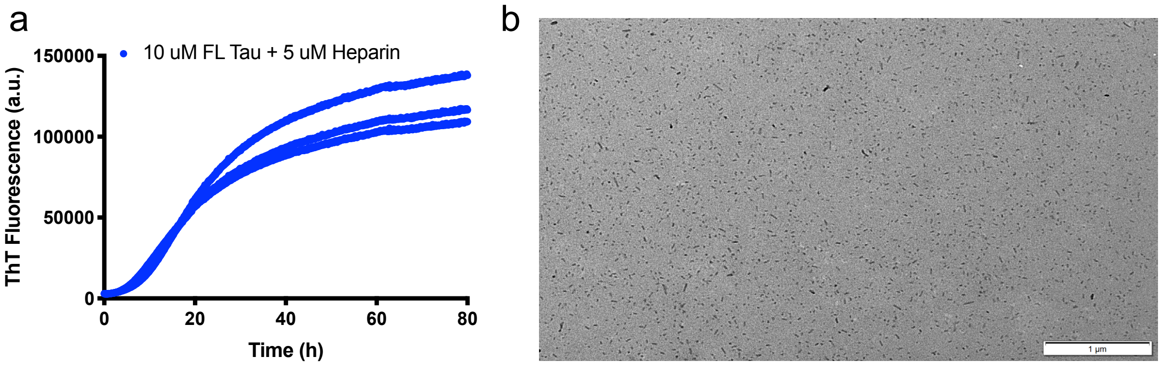


**Supplementary Figure 5. Preparation of recombinant full-length (2N4R) tau seeds.** (a) Th-T kinetics were performed in triplicates to monitor the aggregation of full-length tau$^{2N4R}$ at 10$\mu$M over time. (b) Electron micrographs validate that uniform tau seeds are formed after sonication of end-state tau amyloid fibrils.

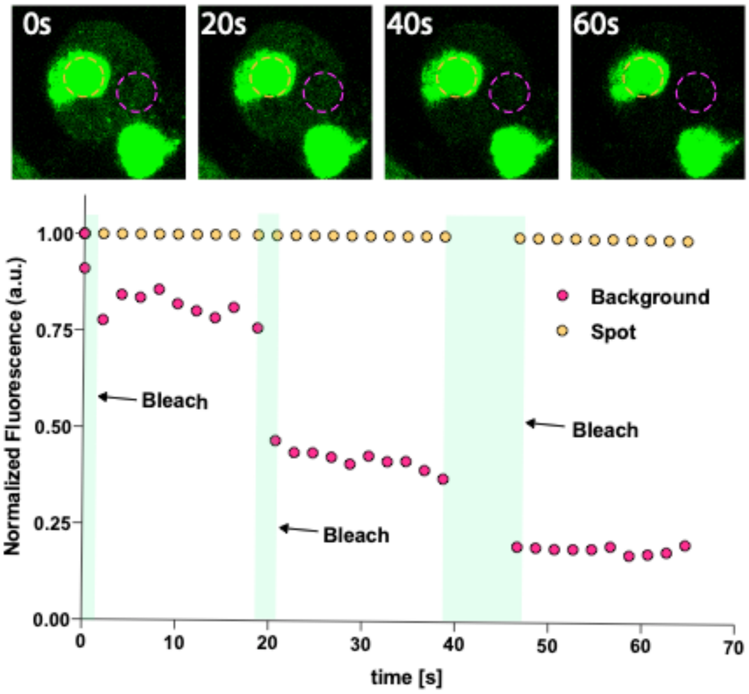


**Supplementary Figure 6. Fluorescence recovery after photobleaching (FRAP) measurements of tau inclusions in the FRET biosensors.** Fluorescence recovery was measured for a region defined within a tau spot (yellow ROI) and a region within the cytoplasm of the cell (magenta ROI), after successive bleaching steps performed in the cytoplasm. Individual timeframes validate that the background of the cell containing soluble tau is bleached, whereas fluorescence of the puncta remains unaffected indicating that there is minimal trade-off with the soluble protein in the cell.

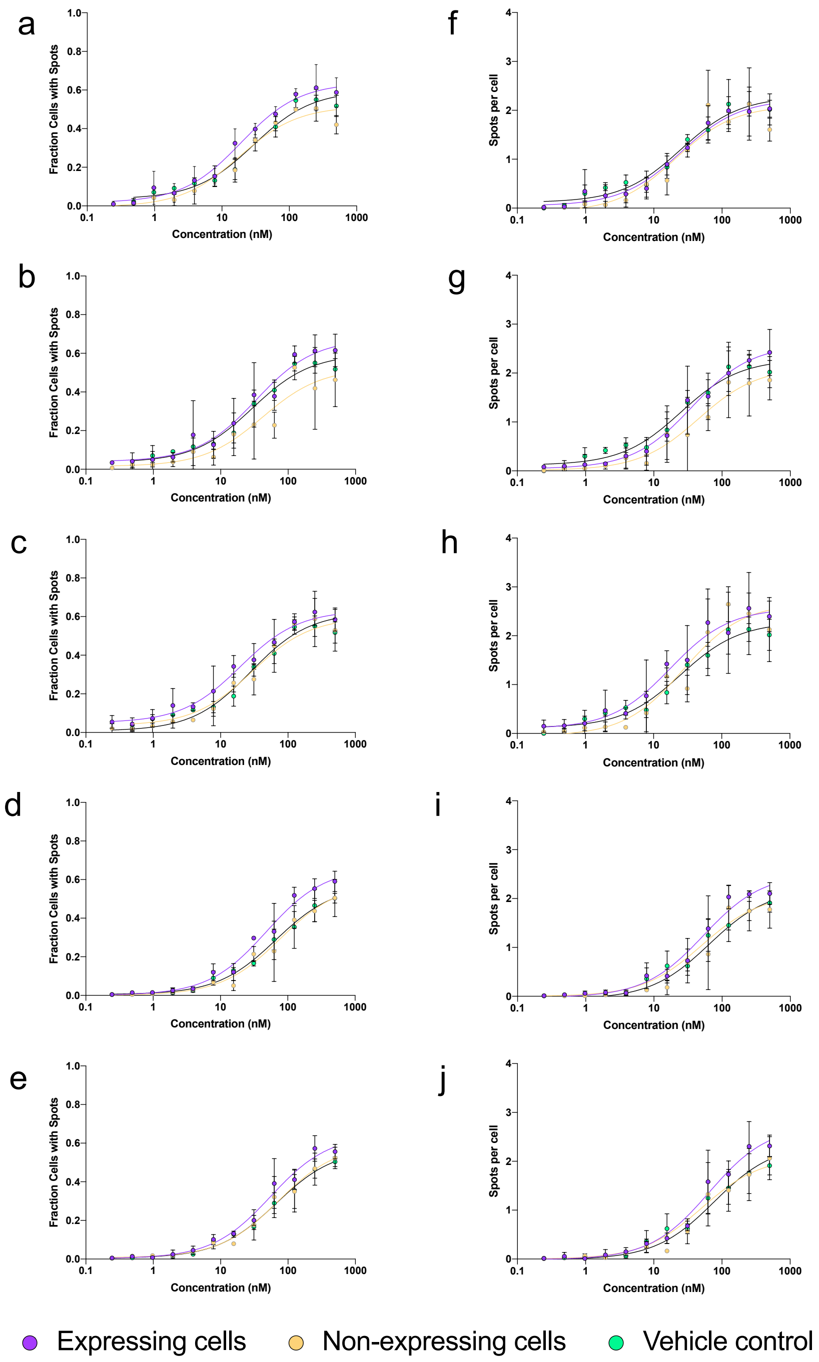


**Supplementary Figure 7. Construct co-expression in the FRET biosensors, following by concentration-dependent tau seeding.** Number of cells with spots (a-e) and number of spots per cell (f-j) that were identified in the FRET cell line for a concentration gradient of tau seeds, following transfection with (a, f) RAB3GAP1, (b, g) CALY, (c, h) NPAS3(d, i) MSR1 and (e, j) TIAM1, respectively. Minimal differences compared to the vehicle and non-expressing cells indicate that expression of these constructs does not affect tau aggregation and spreading. Experiments were performed in triplicates as individual 96-well plate preparations.

### Supplementary References

1 Louros, N., Orlando, G., De Vleeschouwer, M., Rousseau, F. & Schymkowitz, J. Structure-based machine-guided mapping of amyloid sequence space reveals uncharted sequence clusters with higher solubilities. *Nat Commun* **11**, 3314, doi:10.1038/s41467-020-17207-3 (2020).
